# Challenges in Peptide-Spectrum Matching: a Robust and Reproducible Statistical Framework for Removing Low-Accuracy, High-Scoring Hits

**DOI:** 10.1101/839290

**Authors:** Shane L. Hubler, Praveen Kumar, Subina Mehta, Caleb Easterly, James E. Johnson, Pratik D. Jagtap, Timothy J. Griffin

**Affiliations:** Rhapsody Data, LLC, Madison, WI 53705; Department of Department of Biochemistry, Molecular Biology and Biophysics, University of Minnesota, Minneapolis, MN 55455; Minnesota Supercomputing Institute, University of Minnesota, Minneapolis, MN 55455

**Keywords:** Peptide-Spectrum Match, tandem mass spectrometry, precursor mass discrepancy, false discovery rate, statistical analysis, proteogenomics, metaproteomics

## Abstract

Workflows for large-scale (MS)-based shotgun proteomics can potentially lead to costly errors in the form of incorrect peptide spectrum matches (PSMs). To improve robustness of these workflows, we have investigated the use of the precursor mass discrepancy (PMD) to detect and filter potentially false PSMs that have, nonetheless, a high confidence score. We identified and addressed three cases of unexpected bias in PMD results: time of acquisition within a LC-MS run, decoy PSMs, and length of peptide. We created a post-analysis Bayesian confidence measure based on score and PMD, called PMD-FDR. We tested PMD-FDR on four datasets across three types of MS-based proteomics projects: standard (single organism; reference database), proteogenomics (single organism; customized genomic-based database plus reference), and metaproteomics (microorganism community; customized conglomerate database). On a ground truth dataset and other representative data, PMD-FDR was able to detect 60-80% of likely incorrect PSMs (false-hits) while losing only 5% of correct PSMs (true-hits). PMD-FDR can also be used to evaluate data quality for results generated within different experimental PSM-generating workflows, assisting in method development. Going forward, PMD-FDR should provide detection of high-scoring but likely false-hits, aiding applications which rely heavily on accurate PSMs, such as proteogenomics and metaproteomics.

## Introduction

Proteomics is now an important tool in many biological and medical applications. One of the common proteomics tools is shotgun proteomics where mass spectrometry (MS) is used to identify fractionated peptides within complex mixtures, derived from proteolysis of intact proteins from a biological sample. After high-performance liquid chromatography (HPLC), a mass spectrometer generates a series of digital files representing many mass spectra, including tandem mass spectra (MS/MS), which contain information on fragmented peptide sequences. There are many protocols for analyzing these files, but the initial goal of these protocols is to assign a single peptide sequence to each MS/MS spectra in the dataset. Each assignment is referred to as a peptide spectrum match (PSM). In addition, each PSM comes with a score, a measure of confidence that the assignment is correct. The PSMs are then filtered based on score, generating a list of peptides for further analysis, such as inferring the presence of specific proteins within the sample.

Recently, the emergence of new applications such as metaproteomics ^1–2^ and proteogenomics^3–4^ has led to a new emphasis on the accuracy of PSMs^3, 5–7^. These new applications, along with a steady interest in post-translational modification (PTM) identifications depend heavily on peptide-level identifications (Figure 1). For example, metaproteomics researchers might find that a PSM identifies a peptide mapping to a specific species within a complex community of bacteria, and proteogenomics researchers might use a PSM to identify a novel disease-associated peptide carrying an amino acid sequence variation. For all these applications, inaccurate PSMs can support erroneous and costly conclusions, leading researchers to waste resources and time on validating false leads.

**Figure 1.**
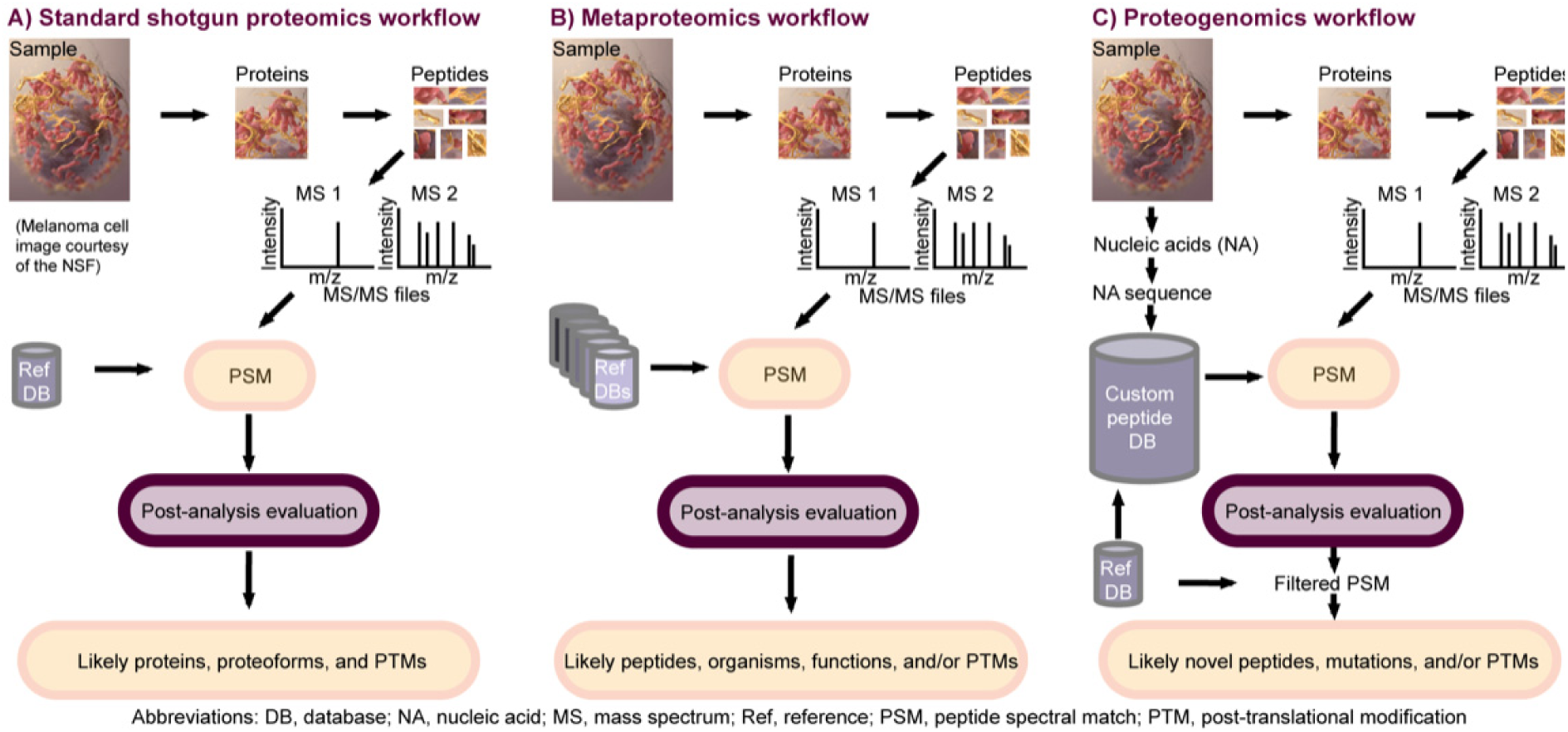
Three common types of MS-based proteomics workflows, all of which depend heavily on accurate PSMs. Standard shotgun proteomics (A) matches MS/MS data against a reference database of known protein sequences, attempting to identify peptides (with and without PTMs), and, finally, infer proteins. In metaproteomics (B), proteins are isolated from a community of microorganisms, analyzed by LC-MS/MS, and matched against an aggregate MS/MS protein sequence database encompassing all probable organisms in the community. These sequences are used to determine taxonomic composition and functional state of the system. In proteogenomics (C), nucleic acid sequence data are obtained from the sample and translated *in silico* to produce a database of hypothetical expressed peptides and proteins, including sequence variants.

Given the central role of accurate PSM assignments to these shotgun proteomics applications, a robust and reproducible statistical framework for determining the probable correctness of PSMs is vital to guide this filtering process to find the peptide sequences most worthy of further investigation^8^. Because it is not possible to classify each putative PSM as true or false with complete certainty, the investigator must accept a defined threshold on a confidence measure, presumably probabilistic in nature, that a particular discovery is likely to be correct. One such statistic is the group-wise false discovery rate (gFDR), defined as the probability that a randomly selected PSM within a group of PSMs is incorrect. Another statistic, the individual FDR (iFDR) (usually called a “local FDR”), often calculated from the gFDR, is the probability that a specific PSM is incorrect. In other words, while gFDR and iFDR ostensibly measure the same thing, iFDR provides a different value for every PSM rather than for the whole group^9^. The greater the accuracy in estimating the gFDR and iFDR statistics, the higher the likelihood of correctly filtering PSMs to those with the highest probability of being correct.

As PSM-centric applications continue to gain momentum, a conceptually simple filter that detects high scoring but likely incorrect PSMs while retaining those of highest confidence would be of great value to many researchers. One metric available for all generated PSM reports, irrespective of algorithm or workflow used, is the precursor mass discrepancy (PMD). The PMD is relative difference between the empirically measured precursor mass and the computed mass of the peptide sequence assigned to the corresponding MS/MS spectrum; this statistic is usually reported in parts-per-million (ppm). A PMD value of zero represents complete concordance between the measured and computed peptide masses; however, PMD can be positive or negative, depending on whether the measured mass is greater or less than the computed mass, respectively.

Traditionally, PMD has been used to *filter* potential candidate peptide sequences for matching to MS/MS^10^, reducing the search space to a more manageable size. Researchers will often use PMD to provide additional evidence (or lack thereof) for a particular PSM – including only PSMs with PMD close to zero for further consideration (e.g. within +/− 5 ppm) after they have been scored by the sequence database search algorithm. Surprisingly, however, PMD is rarely used explicitly to score PSMs.

As such, we sought to improve the accuracy of the gFDR and iFDR statistics using a scalable computational methodology based on PMD that would provide information that is at least partially orthogonal to the existing PSM scoring algorithms. While most algorithms do not use PMD explicitly, there are a few that do, such as Wenger et al. with COMPASS^11^, Cox et al. with Perseus^12^ and Petyuk et al. with DTARefinery^13^ which are focused on shotgun LC-MS/MS data. One method called ProteinProcessor was described for identifying purified proteins from tryptic digests using MALDI-TOF/TOF MS, which included PMD of precursor peptide mass measurements coupled with MS/MS data to increase confident peptide identification and overall protein inference, in a hybrid peptide mass fingerprinting and MS/MS-based method^14^. In addition, some algorithms use PMD and other post-scoring analyses explicitly to change a confidence score (c.f. PeptideProphet^15^, and Percolator^16^), or estimate both iFDR and gFDR^17–19^.

Although these usages of PMD are valuable, here we sought to develop a new and more flexible implementation of PMD, using applications in metaproteomics and proteogenomics as use cases for our work. We conducted a rigorous re-examination of the underlying statistical assumptions traditionally made when employing PMD (e.g. normal distribution of data, time-independence of mass error measures). Our findings demonstrate that many of these assumptions do not hold. As such we have developed a new algorithm that addresses these failed assumptions and that performs post-analysis on previously scored PSMs. Our overarching motivation in this work was to improve confidence in reported PSMs and provide researchers an additional means to assess their accuracy – providing a tool especially valuable for assessing results generated from metaproteomic and proteogenomic pipelines^20–22^.

## Materials and Methods

### Datasets for development and evaluation

To develop and evaluate the PMD-FDR method, we used data from three publicly-available MS-based shotgun proteomics studies ^22–24^, the details of which are shown in Tables 1 and 2. We used four PSM datasets derived from data generated in these three studies. The first of these (*Pyrococcus)* was derived from a standard single-organism proteomics workflow^24^ where PSMs were generated from a *Pyrococcus furiousus* sample, which has a proteome sequence orthogonal to the human proteome (except for only a handful of tryptic peptides). Matching peptide MS/MS spectra from a *Pyrococcus* sample against a protein sequence database that combines sequences from both *Pyrococcus* and humans provides an ideal ground-truth database for testing methods for PSM scoring methods^24–25^. PSMs to human peptide sequences represent false matches, and can be used to estimate actual false positive rates for algorithms being utilized.

**Table 1.** Metaproteomics Dataset Descriptions

|  | Oral 737 (two-step) | Oral 737 (combined) |
| --- | --- | --- |
| **Databases** | Human Oral Microbiome Database (HOMD)<br>Contaminant Database (common repository of adventitious proteins) |  |
| **Manipulations** | Analyzed using a traditional two-step database searching method | Analyzed using a modified two-step database searching method, where the database is searched in sections, and potential proteins present are combined to make a new database which is searched in a second step |
| **Biological Sample** | Cultured oral sample, one participant, without sugar addition |  |
| **Biological Purpose** | Metaproteomics study to investigate effect of sugar on taxonomic diversity of oral biofilms |  |
| **Instrument** | LTQ Orbitrap Velos |  |
| **Link to Publication** | https://microbiomejournal.biomedcentral.com/articles/10.1186/s40168-015-0136-z<br>See reference<sup>23</sup> |  |
| **Link to Public Data** | http://proteomecentral.proteomexchange.org/cgi/GetDataset?ID=PXD003151 |  |

**Table 2.** Single-organism proteomics and proteogenomics dataset descriptions

|  | *Pyrococcus* | Mouse Proteogenomics |
| --- | --- | --- |
| **Databases** | *Pyrococcus furiosus* (Uniprot)<br>Contaminants<br>Human (Uniprot) | Contaminants + Uniprot<br>Mouse: 3-frame translation of Ensemble cDNA<br>Sample RNA-Seq: Observed<br>Splice junction variants<br>3-frame translation of long non-coding RNA (lncRNA)<br>Single amino acid variant (SAV) |
| **Manipulations** | Combined into one database | Combined into one large database<br>Data sectioned into five parts, each matched to MS/MS sequentially<br>Results combined |
| **Biological Sample** | *Pyrococcus furiosus* | Mouse Developmental B-Cell (Pre-Pro-B and Prob-B stages) |
| **Biological Purpose** | Single organism validation data | Prediction of Gene Activity in Early B-cell Development |
| **Instrument** | LTQ Orbitrap Velos | LTQ Orbitrap Elite |
| **Link to Publication** | https://pubs.acs.org/doi/full/10.1021/pr300055q<br>See reference<sup>25</sup> | https://www.ncbi.nlm.nih.gov/pmc/articles/PMC4276347/<br>See reference<sup>22</sup> |
| **Link to Public Data** | https://www.ebi.ac.uk/pride/archive/projects/PXD001077 | https://usegalaxy.org/u/thereddylab/p/prediction-of-gene-activity-in-early-b-cell-development-based-on-an-integrative-multi-omics-analysis |

Two additional datasets included PSMs generated using a metaproteomics analysis of previously generated MS/MS data from human saliva (Oral 737)^23^ using two different workflows implemented in the Galaxy for proteomics (Galaxy-P) platform^5^. One workflow used our previously published two-step method^26^ generating results we called the Oral 737 (two-step) dataset. The second workflow used a more updated approach dividing the large database into smaller sections and matching MS/MS to each section, leading to the creation of a database combining the proteins identified from each section, which are then searched in a second step. We called this the Oral 737 (combined) dataset. These different workflows provided slightly different PSM results from the same input data, and provide valuable datasets for testing the PMD-FDR approach.

A final PSM dataset was derived from a proteogenomics workflow (Mouse Proteogenomics dataset), where transcriptome sequences were used to generate a large database of proteins sequences using data from a previously published study^22^. Collectively, these datasets provided a diverse selection of PSMs from different workflows for testing and evaluating the basis and effectiveness of the PMD-FDR approach. The basic algorithms for sequence database searching and initial PSM score assignments utilized in these workflows to produce these PSM datasets have been described^5, 20^ and are based on the well-described SearchGUI/PeptideShaker platform^27–28^.

### Post-scoring analysis using PMD

#### Qualitative grouping of Peptide-Spectral Matches

In order to perform the necessary post-scoring analysis using PMD, we needed to separate the set of all PSMs into several groups based on peptide length, due to the observed scoring dependence on length. Table 3 describes this methodology and Figure 2 shows the resulting structure for the first dataset (Oral 737 (two-step)). We utilized this data and methodology to test underlying assumptions of PMD filtering and, from the results, developed our modified approach for PSM confidence assessment.

**Figure 2.**
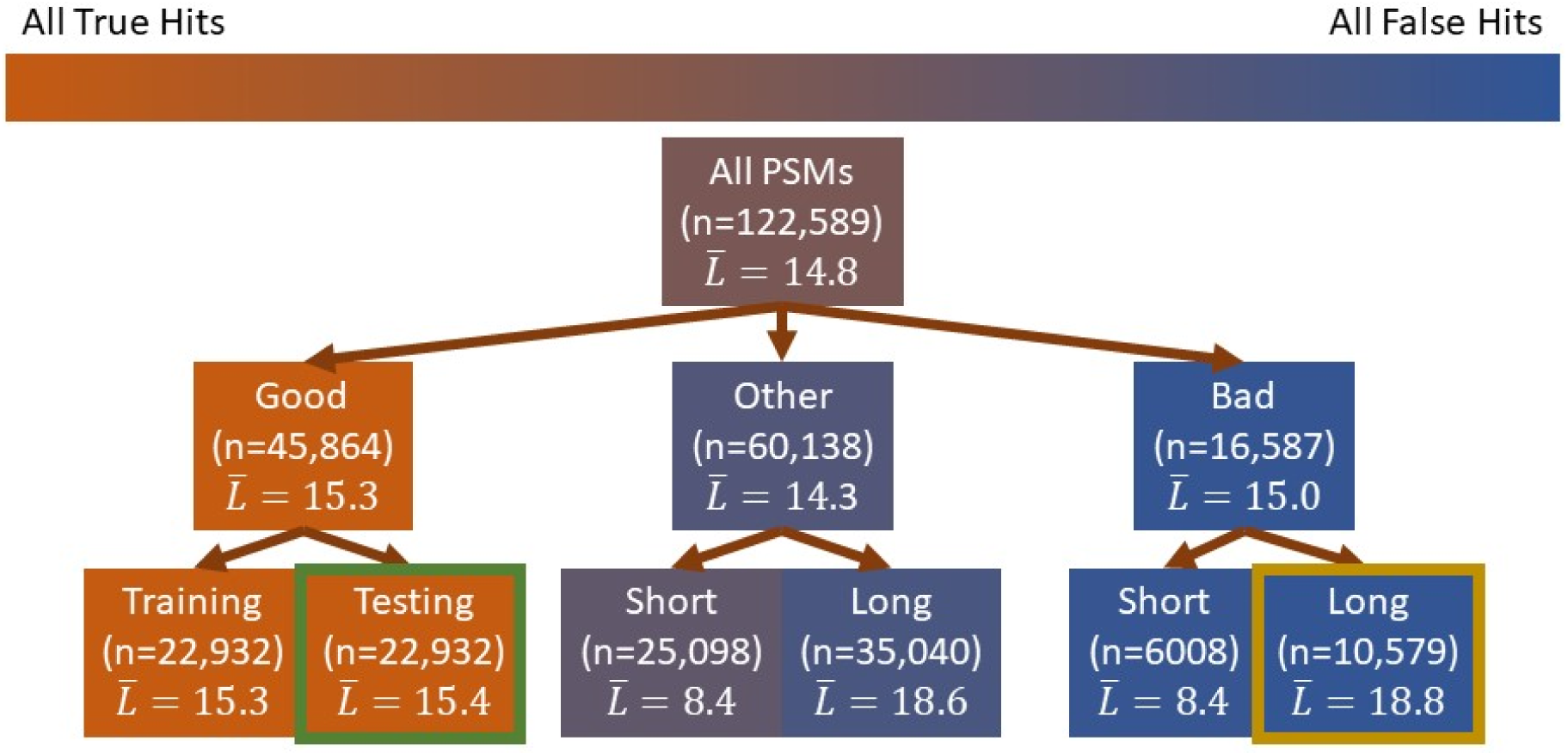
PSM data categories and the numbers within each category for the metaproteomics Oral 737 (two-step) data used for evaluation of our PMD analysis approach. See Table 3 for a definition of the groups. 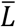 is the average peptide length of the group (number of amino acids). *n* is the size of the group. The green box highlights the group that we used to estimate the density of the correct PSM group (true-hits), while the yellow box highlights the group that we used to estimate the density of the incorrect PSM group (false-hits). Color of the box is a qualitative estimate of the proportion of true-hits in the group.

**Table 3.** Levels of data quality and definitions for PSMs used in this work

| **Term** | **General principle** | **Our implementation** |
| --- | --- | --- |
| True-hit | PSM correctly matches peptide to spectrum | Not implemented. (True-hits are theoretical constructs of which we can never be certain.) |
| False-hit | PSM that incorrectly matches peptide to spectrum | Not implemented. (False-hits are theoretical constructs of which we can rarely be certain.) |
| Good Hit | PSM that is almost certainly a True-hit. | PSMs whose PeptideShaker Confidence score is exactly 100 and the peptide is not a decoy peptide. |
| Bad Hit | A PSM that we are certain is a False-hit | A PSM reporting a match to a decoy peptide |
| Good-Training | The PSM is in the training group (from which we derive local medians), which is a sub-population of the Good PSMs. | Every other PSM found in the Good group (i.e. half of the Good PSMs). |
| Good-Testing | The PSM is in the testing group, the one used to report results. Good-Training and Good-Testing must be disjoint. | Every Good PSM that is not a Good-Training PSM. |
| Bad-Training | A PSM that is used for identifying the PMD distribution of a Bad Hit, one that is unlikely to have a correct precursor mass | Bad Hit PSMs used to define statistics for identifying PSMs most likely false based on PMD |
| Other | Everything that is not Good or Bad. | Everything that is not Good or Bad. |
| Bad-Short | A Bad hit identifying a short peptide. These are more likely to have correct mass than Bad-Testing peptides greater than 11 amino acids | A bad PSM that is less than 11 amino acids long. |
| Bad-Long | A Bad hit that is unlikely to have the correct precursor mass | A bad PSM that is at least 11 amino acids long. These are the PSMs that make up the Bad-Training dataset. |

For evaluating PMD for post-scoring analysis, it was necessary to define populations of PSMs as either “good” or “bad”, to determine whether a PMD analysis could distinguish between these extreme cases. As such, we defined good PSMs as those assigned a perfect confidence score of 100 by the PeptideShaker algorithm, while bad PSMs were matches to decoy peptide sequences which by definition are false. As shown in Table 3, it was then necessary to sub-divide these defined peptides into populations for training the algorithm and then testing out its effectiveness. Ultimately PMD operates on the assumption that any given dataset being analyzes is composed of these two populations – good hits and hits that are false but have still qualified based on set scoring thresholds.

#### Evaluation of common PMD assumptions

We estimated the PMD distribution of true-hits by applying a modified density to the good-testing group, restricting the results to the range of PMD values allowed in the original database search (−10 ppm to +10 ppm). The modified density function assumes that the distribution is unimodal (i.e. a single local maximum in the density plot) and that variation from that assumption is noise. It applies the *density* function from R and rearranges the values on the left of the mode (the maximum point within the unimodal plot) so that they are non-decreasing and rearranged the values on the right of the mode so that they are non-increasing. Further, we applied the same function to the bad group (long peptides only) to estimate the PMD distribution of the false-hits, with false-hits being estimated by PSMs to the reverse-decoy sequences selected by PeptideShaker.

To assess possible PMD effects within false-hits, we evaluated the distribution of PMD for all decoy-matching PSMs. We investigated several potential confounders to using PMD of decoy hits as a proxy for the false-hit distribution: peptide length (number of amino acids), mass, charge state, and isotope state. Of these, we observed that PMD only showed a dependence on peptide length, and none of the other factors. To investigate peptide length, we split the range of lengths (6 – 50) into contiguous groups of approximately equal size, from which we analyzed the bad PSMs (i.e. decoy hits). Once again, we applied the *density* function from R (unmodified this time) to visualize the PMD distributions for each subset of bad PSMs.

To check if there was an edge effect on the distribution of bad PSMs, we needed a statistical test that compared the edge of the density distribution with that of the center. We separated the ppm range (−10 to +10 ppm) into pieces of size 1 ppm and computed the proportion of PSMs falling in that range. Next, we computed a credible interval of the proportion (effectively a Bayesian estimate of the confidence interval), using in-house code to compute the highest posterior density interval to a specified precision (0.001) (method described in **Supplemental Information**). The primary purpose of this experiment was to show that the depletion of the edges of the PMD distribution was statistically significant.

#### Identifying invariants

An important part of creating a successful mixture model is to identify features that can distinguish between multiple groups (here: true- and false-hits). These features must remain constant (invariant) for all members of the specific group; otherwise, one cannot calculate the relative proportion between the two groups. Here, we are estimating the *distributions* of true-hits and false-hits; if we are to use these distributions in a mixture model, they must be invariant throughout the experiment. We can then cross-reference these invariants against the distribution of a mixed population to estimate the proportion of true- and false-hits within that population.

Because our experiments showed that PMD drifts during the acquisition process of LC-MS/MS data (see Results section below and Figure 4), the PMD distribution *is not invariant* for true-hits throughout an MS/MS dataset and needed correction. Using spectrum index (also known as scan number), which is directly related to the time of MS/MS data acquisition within an LC-MS/MS experiment, we sorted the good-training PSMs, and split them into contiguous subsets of approximately 100 good-training elements, being careful not to cross scan number boundaries. Next, we used these temporally ordered subsets to compute a local median, by which we translated *all* PSMs within the same spectrum index range. In other words, we shifted every PSM within a temporal subset by the local median PMD of the training PSMs. Finally, we used good-*testing* PSMs, split into 100 different subsets based on spectrum index of roughly the same size (in terms of good-testing PSMs, not in terms of total PSMs). We were careful to not use the same data (testing vs. training) nor the same data divisions (100 subsets vs. 100 good-training PSMs *per* subset), to avoid introducing analytical artifacts.

#### Score-based FDR (sFDR)

We used the following procedure to assign a PMD-FDR for a particular PSM. We refer to the PMD-FDR of the *i*th PSM as *FDR_i_*. We also need to require that every PSM belongs to some subset of all PSMs; we use *j* to designate the index of that group. PSMs were assembled into groups based on assigned scores, using score ranges to ensure PSMs were divided up into 10 groups of approximately the same size. Analyzing groups of approximately equal size was import to keep similar variance in measured variables between the groups. In order to use Bayes’ Rule to compute an individual FDR from a mixture we need to know three things:

- The likelihood that a true-hit could produce the *i*th PMD, given that it was a true-hit. Designate this *t_i_*.
- The likelihood that a false-hit could produce the *i*th PMD, given that it was a false-hit. Designate this *f_i_*.
- The probability that the PSM is a false-hit, given that the PSM is a member group *j*. Designate this *α_j_*.

Given these requirements and notation, we have the following formula:

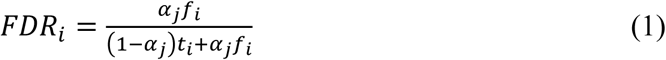

Equation 1 presents the challenge of requiring some form of a group-wise FDR (*α_j_*) to compute the individual FDR. We bootstrap that process by 1) estimating a group-wise FDR for a collection of PSMs with similar scores; 2) we estimate the mean score for each group of scores; and 3) interpolating those scores and corresponding FDR estimates.

The first part: *α_j_* is an estimated FDR for all PSMs with a score in the *j*^th^ range, calculated using

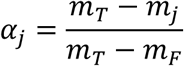

where *m_T_* is the maximum density for the true-hits, *m_j_* is the maximum density for the group *j*, *m_F_* is the maximum density for the false-hits.

Second, we create a function calculating

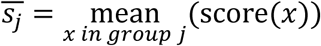

Third, we interpolate the estimated FDR between all pairs 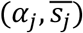, creating a new, interpolation function mapping all possible score values (*S*) to the interval [0,1]

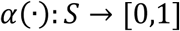

This new value is effectively a groupwise FDR that is specific to any particular PSM based only on its score. It is conceptually similar to grouping all PSMs by score and computing the groupwise-FDR for that subset. However, by using larger groupings of scores, we improve our precision and, hopefully, reduce noise of the measurement. The interpolation creates a more continuous function.

#### Individual FDR (iFDR)

Now we have the tools to estimate iFDR (more commonly known as the local FDR). One of the issues that arises when using the formula above to estimate *FDR_i_* is that we must have a large enough population in group *j* to be able to estimate *α_j_*. However, by creating a score-based interpolation, we can use the entire dataset to estimate the group-wise FDR for a given score. In other words, by replacing *α_j_* by *α*(*s_i_*) in the formula for *FDR_i_*(above) we get the following estimate of the *i*^th^ FDR, without explicit reference to the subsets of the data.

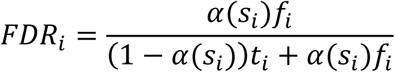

#### Groupwise FDR for arbitrary subsets (gFDR)

The gFDR represents the overall FDR for an arbitrary group of PSMs. Starting with the iFDR values, the gFDR is simply the mean iFDR across the entire group.

#### Illustrating effectiveness of PMD-FDR

To show that PMD-FDR can assist in separating good hits from bad when addressing high-scoring PSMs, we used a publicly available non-human dataset (Pyrococcus_tr) that was analyzed using a two-species reference database that combined *Pyrococcus* and human reference databases, where the human reference database was used as a relatively large confounder containing sequences known not to be present within the *Pyrococcus* sample analyzed. Adding this confounding database mimics the situation encountered in metaproteomic and proteogenomic applications, where the database contains a relatively small number of protein sequences actually present in the sample, along with many sequences which are essentially noise. Note that a decoy database was also appended, in the form of a reversed sequence database created from the entire two-species reference. We calculated PMD-FDR for all of the PSMs. Next, for every confidence score and group (human, *Pyrococcus*, contaminant, and decoy) we computed the rate of rejection; that is, given a confidence score, *C*, the proportion of PSMs in the subset with a score above *C* that had a PMD-FDR greater than 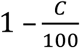. We performed this calculation for all integer values *C* from 1 to 100 and for all four groups.

## Results

### PMD analysis of PSMs

We first tested the implicit assumption that the PMD values for a group of true identifications will have a bell-shaped distribution (or, more strongly, a normal distribution) centered on zero, which can be approximated by restricting the analysis to the highest-scoring PSMs (Figure 3, black line). Unsurprisingly, while the distribution for this group is usually (but not always) bell-shaped, it is slightly off-centered – a result of systematic measurement error in the instrument. We observed this effect to varying degrees in every dataset we analyzed (see **Supplemental Information**), indicating that the assumption of a distribution centered on zero does not always hold. If we are to use PMD to estimate FDR, we must allow for this shift in mass discrepancy, regardless of the direction of that shift. We call this systematic shift the PMD-Shift, regardless of cause.

**Figure 3.**
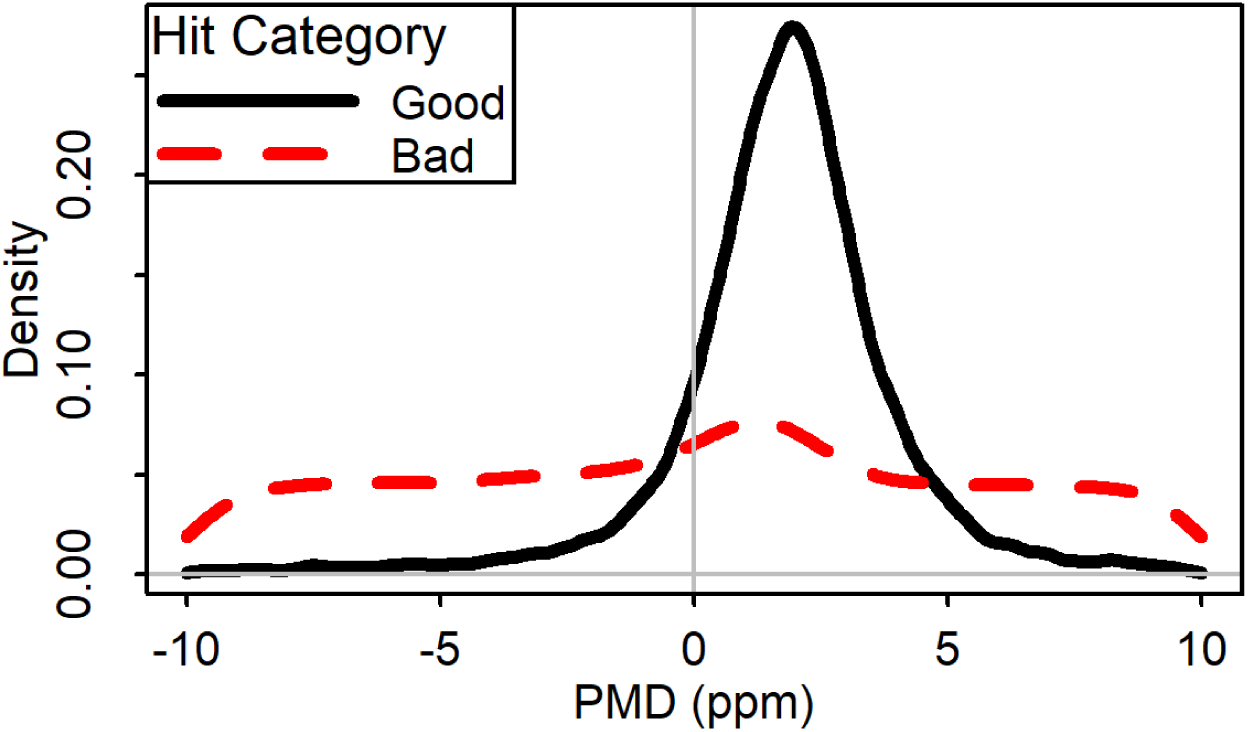
PMD values for good and bad peptide identifications in the Oral 737 metaproteomics (two-step) data set. Shown is a density plot where the area under the curve totals 1. The solid line represents PSMs from the Good-Testing group. The dashed line represents PSMs from the Bad group (both short and long). This shows a systematic shift of accuracy for Good hits (PMD-shift) and a slight preference for accuracy among decoys (the Decoy-Mode).

Concerning the distribution of false-hits, there are typically two similar but distinct assumptions (c.f. ^17^): that they are either uniformly distributed or normally distributed with large variance. We found, however, that plotting the density of decoy hits yielded mostly, but not completely, uniform distribution with a rise close to zero (Figure 3, dotted red line). Once again, the assumptions did not hold; this also held true for each of our four datasets. Interestingly, the mode of the decoys is roughly the same as the mode of the good hits. We will refer to this effect as the Decoy-Mode.

A more subtle assumption used in any mixture model, one that is so fundamental it is rarely mentioned, is that the probability distribution of each group (e.g. true-hits and false-hits) across the dependent variable (here, PMD) does not change during data acquisition; i.e., the mixture model assumes that PMD distribution is “invariant” across the time period of the LC-MS/MS run. We found this assumption is also false (Figure 4a) – the accuracy of the PMD *can* change during an experiment. Fortunately, we have identified a single variable, spectrum index (scan number concatenated with file name), that directly corresponds to the time of acquisition for any MS/MS spectrum, *and* is highly correlated with the PMD-Shift. Thus, after subtracting the local medians from the good-training group, we have a much more consistent accuracy, centered about zero, with similar variance throughout the dataset (Figure 4b).

**Figure 4.**
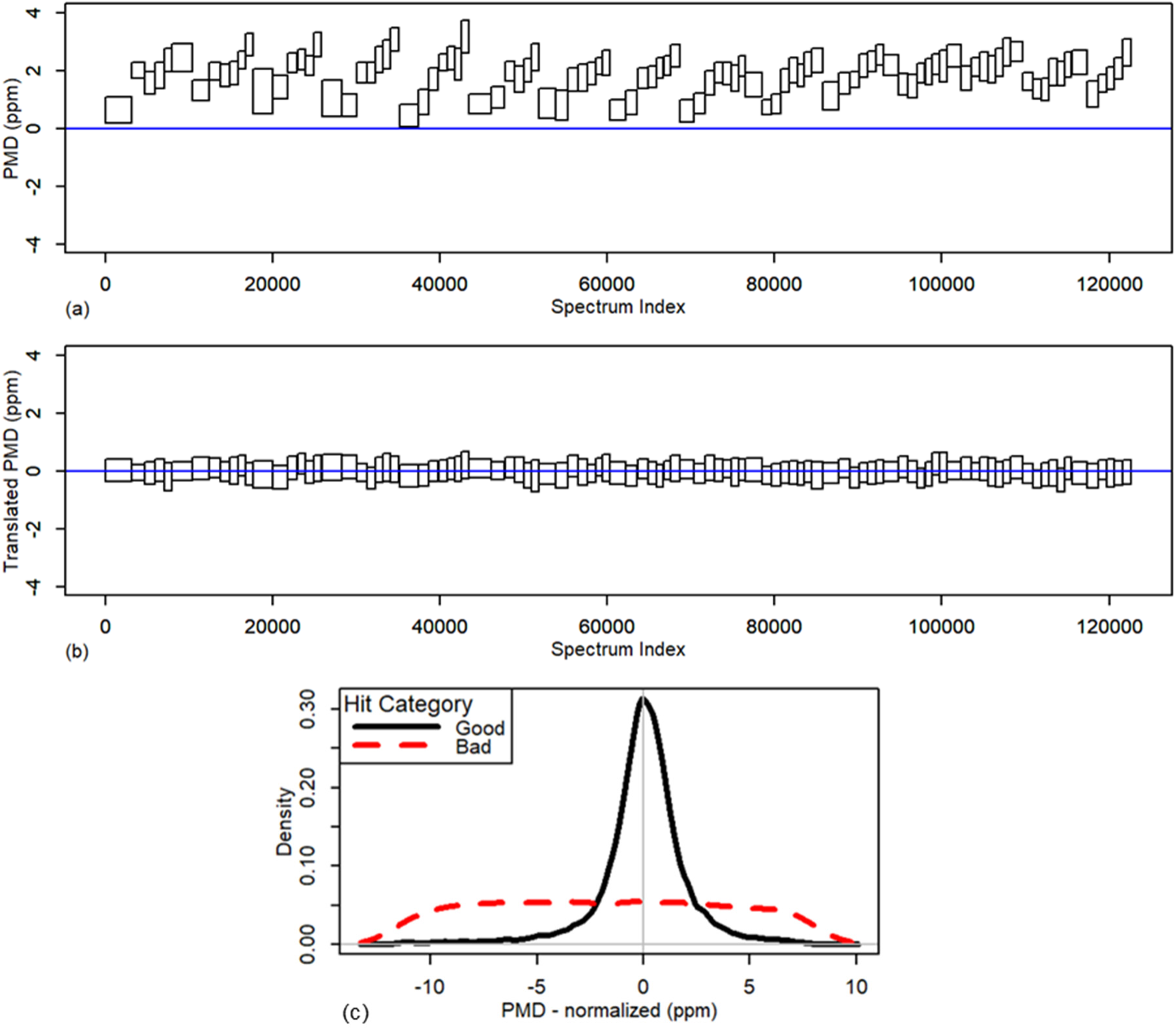
Middle 25% of PMD values A) before and B) after translation for subsets in the Oral 737 metaproteomics data set plotted as a function of spectrum index number (a numeric assignment that combines spectrum number and spectrum file name). Note that each box represents 1% of the spectra in the good-testing group; the number of items in the group overall changes based on the density of the good-testing PSMs within that group. C) Oral 737 results from Figure 3 above after translation using PMD correction.

In order to identify the probable cause of the Decoy-Mode (shown in Figure 3), we plotted the distribution of the decoys, conditioning on a variety of quality metrics. Figure 5 shows results when plotting the PMD distributions by length of predicted peptide. Of particular interest and relevance is that peptides of length 6-8 amino acids had a strong Decoy-Mode, whereas the groups whose peptides were all longer than 12 amino acids were completely free of the Decoy-Mode.

**Figure 5.**
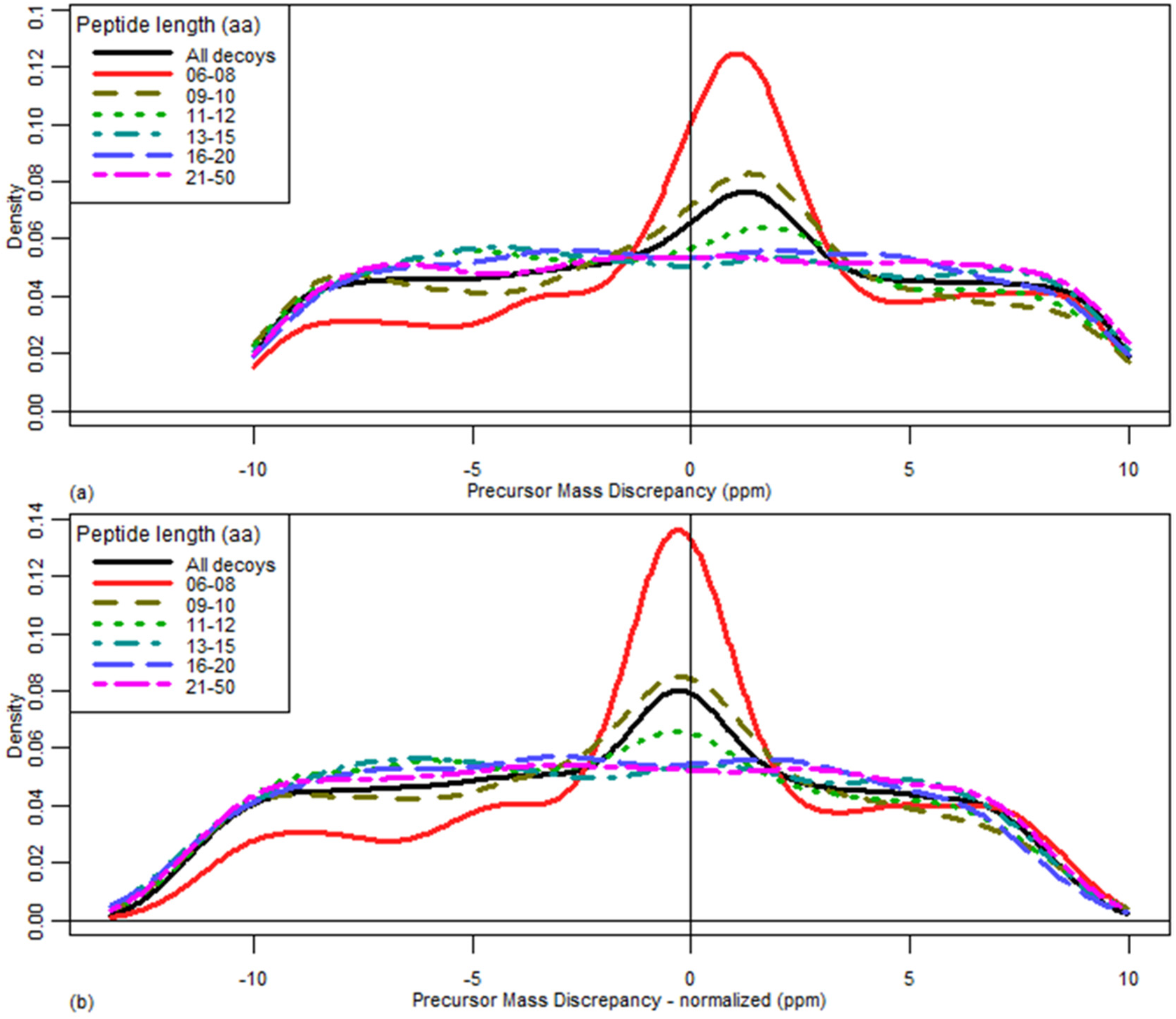
Decoy-Mode: A) Distribution of PMD for decoy (bad) PSMs, conditioned on the number of amino acids in the identified peptide. B) PMD distribution for decoy (bad) PSMs condition on the amino acid length after translation for the PMD shift as described above in Figure 4.

Finally, we observed the decoy distribution also appears to decrease near the edges of the PMD range (Figure 3). Since this could potentially be an edge-effect artifact of using the R package^29^ *density* function, we decided to compare the proportion of decoys in PMD values in each ppm range (−10 ppm to −9 ppm, −9 ppm to −8 ppm, etc.) to see if the shape of the decoys changes significantly (Figure 6). In fact, the probability of extreme values is significantly less than for non-extreme values; in particular, the most extreme 4 ppm each have a proportion that is significantly less than the overall proportion. This suggests that using a uniform distribution to estimate the distribution of false-hits may be misleading for extreme values. The normal distribution assumption can likewise be discounted because of the relative *uniformity* of the plot. Here we will refer to this effect as Decoy-Tail.

**Figure 6.**
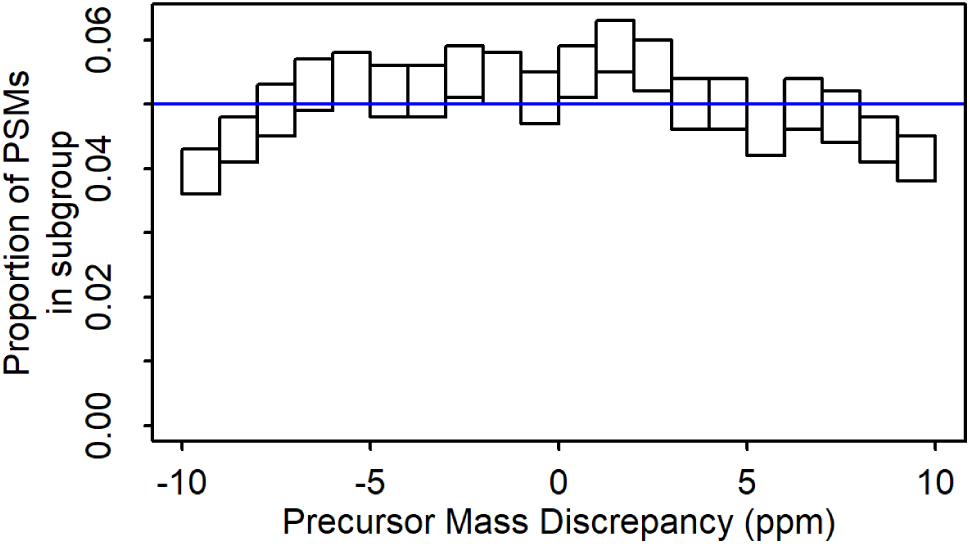
Proportion of decoy (bad) PSMs by PMD. We split the range of PMD values into twenty equal parts and computed the proportion of decoy PSMs within that range. The width of the boxes reflects the range of PMD values considered for a group. The height represents a 95% credible interval for the proportion of PMD values in this region; this interval is the highest posterior density interval (see Supplementary Information for more details). If the distribution of PMDs for decoy PSMs was uniform we would expect that roughly 95% of all of these intervals include the blue line (i.e. approximately two exceptions) and, in addition, that the exceptions would not have a strong spatial pattern.

**Figure 7.**
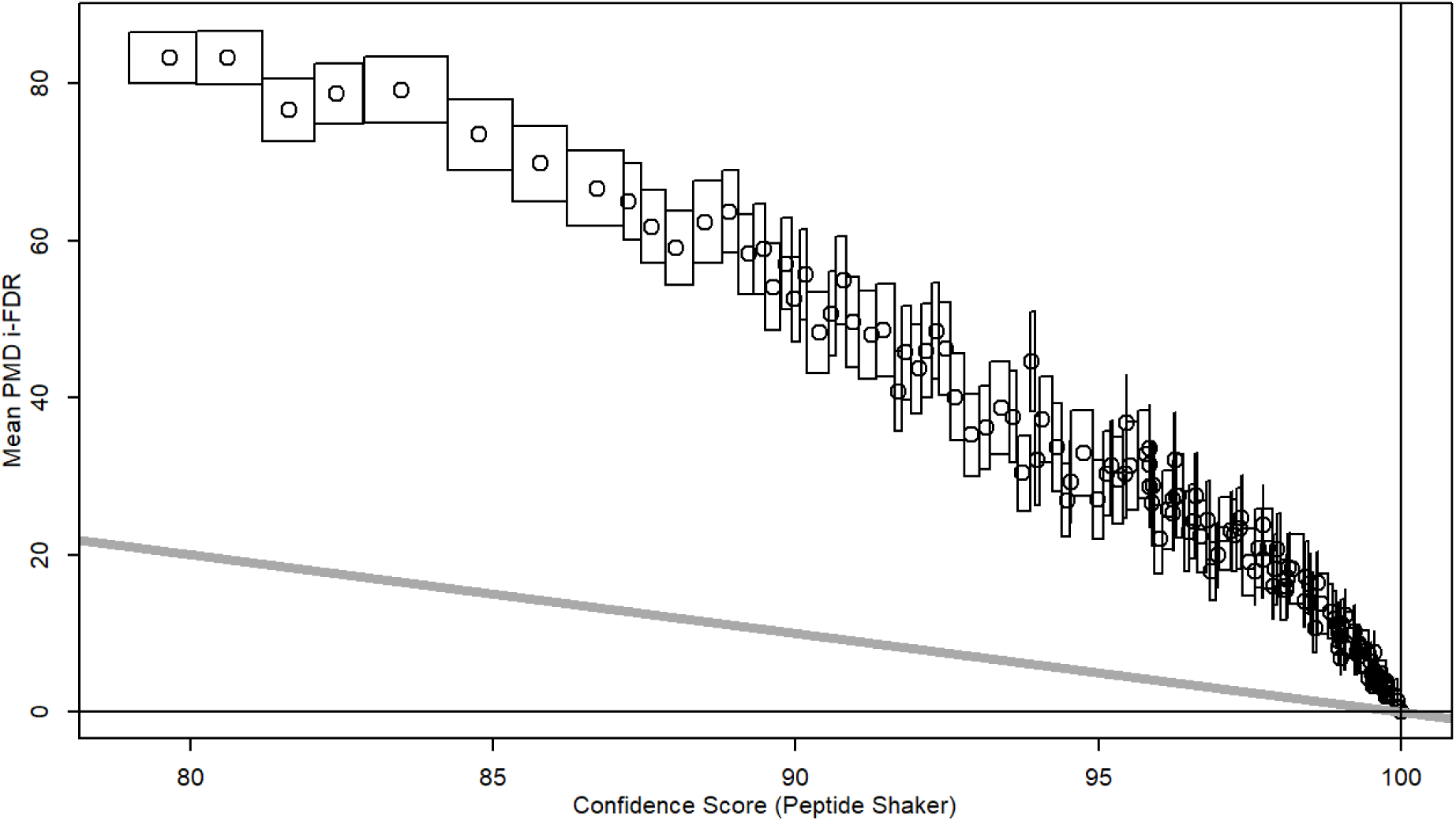
Comparison of PMD-FDR, Target-decoy-FDR (TD-FDR), and Confidence (PeptideShaker). All (and only) PSMs reporting 1% TD-FDR are included in this chart. Each box represents 100 PSMs, grouped by Confidence score; the width covers the range of Confidence scores while the height is 95% confidence interval of the mean PMD-FDR. The circle represents the mean Confidence and PMD-FDR. The gray line is a reference for perfect agreement between PMD-FDR and Confidence (i.e. here they only agree where they must, by definition).

These three issues, the Decoy-Mode, PMD-Shift, and the Decoy-Tail, can cause a mixture model using PMD to fail; that is, produce an inaccurate FDR. However, if we address these issues then the simple requirements of the mixture model can be upheld.

### PMD translation to generate invariant distributions of PSMs

In order to use a mixture model of PMD to model the combination of true and false PSMs, we need to provide a transformation of PMD that produces distributions that are invariant across the experiment. In the previous section we showed that the raw PMD is not up to the task: the true-hit distribution of PMD can vary significantly, even within a single run, while the false-hit distribution changes depending on the amino acid length of the identified peptide and is neither uniformly- nor normally-distributed. This is the reason for translating the PMD and removing short peptides from consideration, at least when estimating the distributions of true- and false-hits.

Once we have estimated the true- and false-PSM distributions of the translated data, we need to show that these distributions are invariant across the time coordinate of the LC-MS/MS run. Ideally, we would be able to consider every possible confounder. However, for practical reasons, we restricted ourselves here to the two variables that were confounded with PMD: time of acquisition (i.e. “scan index”) and peptide length.

Figure 4b shows the effect of PMD-translation on good hits: as expected, it shifts the mean to (approximately) zero but, more importantly, the spread of good hits (i.e. precision) is also consistent across the range of spectrum indices – the time-dependent PMD-shift has been addressed with no ill effects. Additionally, when using PMD-translation and accounting for peptide length, the distribution of PSMs for decoy peptides longer than 10 amino acids no longer shows a length dependence (Figure 5) -- in other words, by removing peptides < 10 amino acids we now have an estimate of the false-hit distribution that is independent of the peptide length.

### Comparison of false discovery rate (FDR) methodologies

With a working method for measuring PMD and associated statistics, we next sought to compare confidence using traditional PSM scoring (in this case PeptideShaker Confidence scores) to that offered by post-scoring PMD False Discovery Rate analysis (PMD-FDR). Such a comparison is warranted as a step towards understanding how PMD-FDR might be useful as a post-scoring analysis method. PSM confidence derived from PeptideShaker’s Confidence score and PMD-FDR are estimating complementary statistics (see ^28^, and Supplemental Information, Section 4.0 for a description of how Confidence is calculated). Additionally, Table 4 compares these two metrics for two different datasets and for four levels of certainty.

**Table 4.** Comparison of PMD-FDR with PeptideShaker’s Confidence score

| Dataset | PeptideShaker<br>Confidence<br>Score Range | Number<br>of PSMs | PeptideShaker<br>gFDR<br>(100 – mean<br>Confidence)<br>(%) | PMD<br>gFDR<br>(%) |
| --- | --- | --- | --- | --- |
| Oral 737<br>(two-step) | 100 | 45,864 | * 0.0 | * 0.0 |
|  | 99-100 | 5,116 | 0.4 | 4.8 |
|  | 90-99 | 9,180 | 4.4 | 29.2 |
|  | All | 122,589 | 18.7 | 50.6 |
| *Pyrococcus*<br>(using<br>combined<br>*Pyrococcus*-<br>human<br>sequence<br>database) | 100 | 9,243 | * 0.0 | * 0.0 |
|  | 99-100 | 128 | 0.7 | 22.7 |
|  | 90-99 | 746 | 3.4 | 26.5 |
|  | All | 15,111 | 29.9 | 32.5 |

\* By definition.

By far the most common confidence measure of a PSM is the local FDR calculated from a target-decoy (TD-FDR) methodology; standard practice reports PSMs that have 1% TD-FDR ^30^. Figure shows the relationship between PMD-FDR, PeptideShaker’s Confidence score, and 1% TD-FDR. First, there is a perfect agreement when the Confidence score is 100. This is by definition: we used this score to define Good Hits and no decoys can have a perfect score of 100. Furthermore, a large proportion of high confidence PSMs in this group also have a low PMD-FDR (i.e. the two scores are concordant for the highest of the high-scoring PSMs).

However, as the confidence score approaches the lowest in this group (Confidence = 80), PMD-FDR soars to an *average* of 80%. In other words, this group has a low probability of a decoy (1% TD-FDR), PeptideShaker declares it to be 80% likely to be a *true*-hit (Confidence ≈ 80), but PMD-FDR reports that a member of this group is 80% likely to be a *false-*hit. Indeed, by counting squares in the figure, we can observe that there are approximately 2000 PSMs with Confidence between 80 and 90 with 1% TD-FDR, whose overall PMD-FDR is well over 50%; these three PSM confidence measures are related but do not agree on the specifics.

### Detection of False-hits using PMD-FDR

To show that PMD-FDR helps detect, and if desired, remove high-scoring, but false-hits, we applied our methodology to the *Pyrococcus* dataset. To review, this dataset involved the analysis of a *Pyrococcus* data sample using both *Pyrococcus* and human reference databases, and has been suggested as an ideal dataset for testing peptide spectral matching algorithms ^24–25^. PSMs in this dataset are derived from a *Pyrococcus* sample, which has a proteome sequence orthogonal to the human proteome (except for only a handful of tryptic peptides), such that hits to human sequences can easily be assigned as false. As such *Pyrococcus* combined with human protein sequences has been proposed as ideal ground-truth database for testing methods for PSM scoring methods^24–25^. Our goal here was to show that PMD-FDR selectively reduced our confidence in human peptide identifications, which should not be present in the sample (we treated potential contaminants, such as human keratin, as a separate type of identification, to which we assigned the label “contaminant”).

Figure 8 plots the rejection rate of PSMs using any Confidence score (from 1 to 100) as the threshold. This plot is based on results from PSM analysis of *Pyrococcus* MS/MS data against the combined *Pyrococcus*-Human protein sequence database, which can be considered a “ground truth” dataset – hits to *Pyrococcus* can be considered as good matches while those matching human sequences are false. Inspection of the plot provides interesting insights. For example, the proportion of human PSMs with Confidence over 40 that have a PMD-FDR greater than 60% (i.e. a PMD-based confidence less than 40%) is approximately 70%. Note that the rejection rate for human and decoy datasets are approximately constant and equal across the entire range of confidence scores (70-80%) while the rate for “contaminant” and *Pyrococcus* is always less than 10%. The reduction in rejection rates for Human at Confidence = 100 is an artifact caused by our assumption that we consider all PSMs with a Confidence score of 100 to be correct; there were 3 such human PSMs (implying that the actual FDR for this group is greater than 0, though not large).

**Figure 8.**
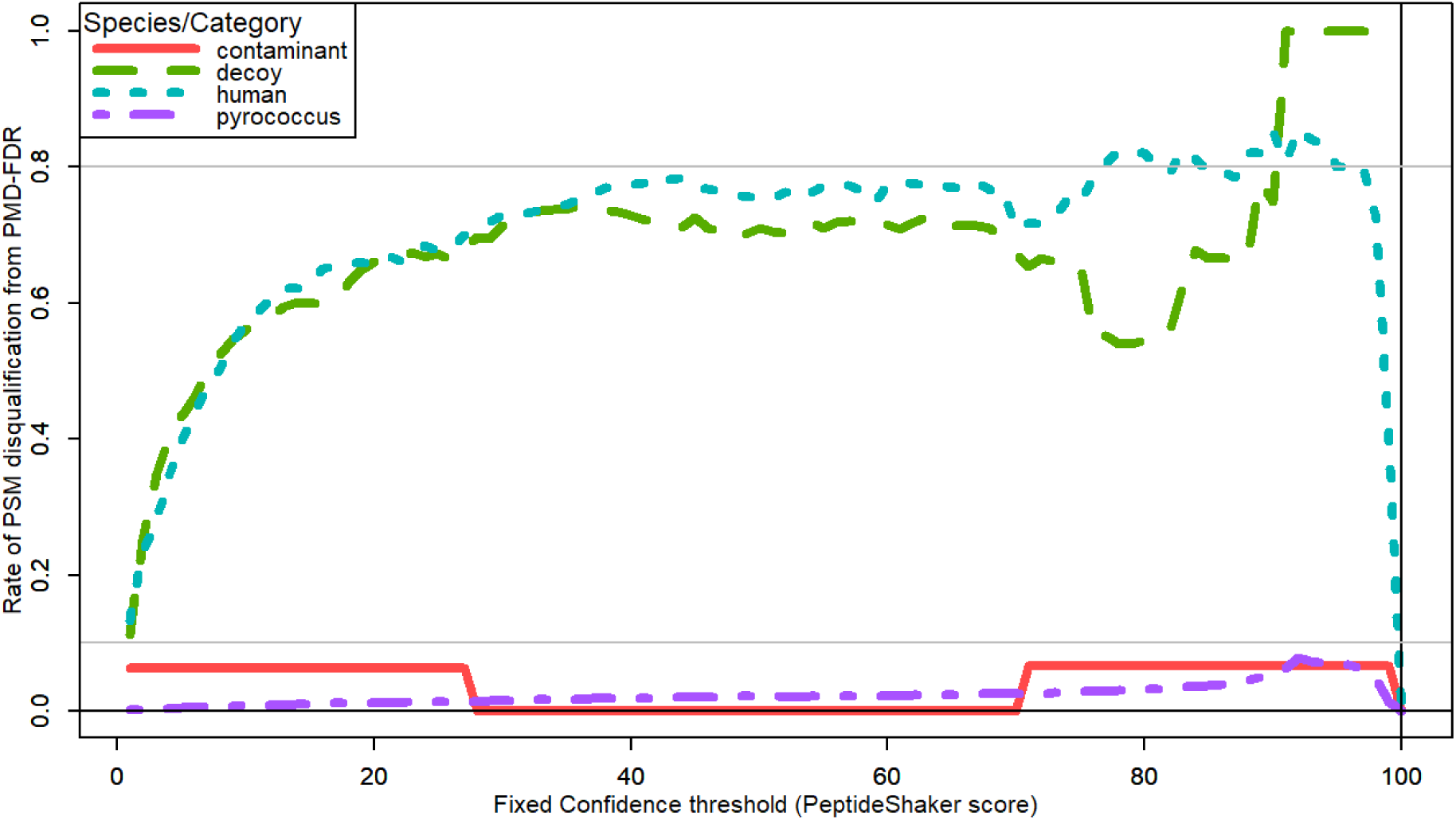
PMD-FDR detects and provides a means to remove likely false-hits when it is used as the threshold. Data was obtained by matching MS/MS data to the combined *Pyrococcus*-Human protein sequence database. Human proteins represent false-hits. “Contaminant” matches to known contaminant proteins, often introduced by sample processing – this includes several human proteins and the “contaminant” designation takes precedence over the “human” designation. Each line represents the PMD rejection (disqualification) rate for a class of peptide: for every Confidence value C, we compute the ratio of PSMs rejected because the PMD-FDR is greater than (100-C)/100, given that the PSM had a Confidence greater than C.

### Application to datasets

In Supporting Information we include analyses of all of the example analytical datasets (See Figure 1 and Methods section for descriptions), derived from three different proteomic studies. We have not shown results from these in the main text due to space constraints, but instead we provide a brief summary:

- We included several different types of representative analyses: metaproteomics data analyzed using two different workflows (Figure S-1 and S-2), single-organisms proteomics (Figure S-3) and a proteogenomics dataset (Figure S-4). These types of projects represent the types of experimental datasets where the PMD-FDR method should have value.
- Two different metaproteomics workflows applied to the same raw data (Figures S-1 and S-2) resulted in very similar true- and false-hit distributions. This means that the PMD-FDR of individual PSMs would be approximately the same between the different analyses. This result is reassuring – the measurement is consistent with the assumption that the PMD distribution for true-hits does not depend on how we generate the PSMs.
- The Decoy-Mode, PMD-Shift, and Decoy-Tail were all present, to a greater or lesser extent, in all four analyses (Figures S-1, S-2, S-3 and S-4).
- The PMD-Shift varied greatly between different datasets – for some it was no more than 2-ppm while for others the data shifted by more than 10-PPM. Note that if someone were to analyze this last case using a small window, say +/− 5 ppm, they would have lost a significant number of good PSMs. Accounting for this shift allows application of the PMD-FDR method to data acquired using different instrumental conditions.

## Discussion

Our PMD analysis of a number of datasets revealed a number of findings, some supporting previous findings, while others provided novel results challenging inherent assumptions used in scoring PSMs. For example, our results confirmed findings of others^11–13^, that the PMD of good hits are dependent on time of acquisition during an LC-MS/MS run, which we called the PMD-Shift. This shift must be accounted for in an assessment of confidence of PSMs. We also identified two structural elements to the PMD distribution of decoys: the Decoy-Mode (a prevalence for small peptides to have more accurate precursor masses) and the Decoy-Tail (a bias for decoy PSMs away from the edges of the precursor range). In particular, these imply that both uniform and normal distributions are poor approximations of the decoy PMD distribution and, presumably, of the false-hit distribution.

Collectively these are novel and significant findings, in that they invalidate the primary assumption of a statistical mixture model: that the probability distributions of the components (here, true- and false-hits) are constant throughout the acquisition of the dataset. Unless one allows for the time-dependence of the PMD of true-hits, the distribution of true-hits is ill-defined. Similarly, unless one allows for the size of the peptide, the PMD of false-hits is ill-defined. To allow for these issues, we removed small peptides when approximating the false-hit distribution and shifted all PMD calculations by a local median of “good” hits.

After performing these transformations, we verified that the resulting statistics were invariant and we created a function that translates scores into an estimated FDR for PSMs with that score. Combining this score with the revised mixture model gives us a local PMD-FDR for each PSM (a posterior error probability (PEP) for a specific type of error: an incorrect precursor mass). Now we can calculate the PMD-FDR for any group of PSMs by simply averaging all the probabilities for elements of that group together. This provides for a practical and useful tool for researchers seeking to further assess confidence in high scoring PSMs. We can use this new score as a filter on previously accepted PSMs to improve our confidence in those that are most reliable. This is valuable for researchers seeking to distinguish high-scoring PSMs of highest confidence from those which should be approached with skepticism – particularly useful for PSM-centric applications such as proteogenomics and metaproteomics. The addition of the PMD-FDR scoring provides added-value to these applications, empowering the individual researcher with additional information on the confidence of PSMs such that they can make decisions on which of these to further validate and which to potentially ignore.

In general, we found PMD-FDR to be more restrictive than either the standard target-decoy FDR or the Confidence score derived by PeptideShaker^28^, which is a score compiled from multiple scoring algorithms employed by SearchGUI^27^. While we used this score to evaluate our methodology because of its apparent high quality, it may still be overly optimistic. Notably, our PMD-FDR method is applied after scoring of PSMs by the database search program, and could be used on results from any upstream program that generates and scores PSMs.

Furthermore, and highly relevant to workflows producing large datasets of PSMs, PMD-FDR can be used to detect biases in methods which may be introducing high scoring false-hits. This can be useful in developing and optimizing new workflows for generating PSMs. Specifically, if we find that two workflows employing different algorithms or parameters produce different numbers of PSMs with 1% target-decoy FDR, then we can use PMD-FDR to determine if we have, in fact, added valuable PSMs rather than adding false-hits with inflated scores.

### Requirements for PMD functionality

In developing the PMD-FDR method, we originally hypothesized that we could provide a largely independent FDR measurement by applying a Bayesian mixture model, assuming that the PMD of true-hits was normally distributed about zero and that the PMD of false-hits were uniformly distributed within the mass measurement error window. In order to use the mixture model, we needed to be able to estimate the distributions of true- and false-hits and show that these distributions are fixed throughout a given experiment. These assumptions did not hold, as PMD distribution for true hits had a complex relationship with time of acquisition (Figure 4a), and the false-hit distribution was dependent on peptide length (Figure 5)

The good news, however, was that both of these difficulties can be overcome by 1) subtracting the local median of the good hits from every PMD ; and 2) restricting our estimation of false-hits to long peptides, say, longer than 10 amino acids. This allows us to identify the distribution of true- and false-hits on a modified PMD score, distributions that are invariant throughout the experiment.

It is also important to note that we found that the false-hit distribution is not well approximated by either a normal distribution (with large variance) nor with a uniform distribution. We suggest that an empirical distribution should be used to estimate both the true- and the false-hit distributions, although it may be reasonable to smooth the resulting distributions, as we have.

### Concordance/Discordance with other confidence measures

When evaluating the degree of concordance between PMD-FDR and the other two confidence measures (TD-FDR and the Confidence score from PeptideShaker) we found concordance with the very high scores – which, in this case is by definition. However, PMD-FDR was much more critical of lower quality spectra than the other two confidence measures – a large number of PSMs reported to be in the 1% TD-FDR group had PMD-FDR of 50% - 100%. Similarly, we reported a 10-fold disparity between PeptideShaker and PMD-FDR for Confidence scores in the 90’s (PeptideShaker effectively reported a 1%-10% FDR where PMD-FDR reported 30%). In other words, we have created a score that agrees with other scores when things are good but encourages greater skepticism when quality drops. As inferred above, we conclude that this implies an independence between PMD-FDR and both the algorithm-specific confidence score (e.g. PeptideShaker’s Confidence score used in this work) and the traditional TD-FDR. We believe that this independence will hold against other algorithms, especially for the many which do not explicitly use PMD in their scoring. Thus, PMD-FDR should make a good, independent filter; in our experiments we observed a reduction in false-hits by 60-80% while only decreasing the number of good hits by 0-7% (Figure 8). Interestingly, this observation is largely independent of score threshold.

### Limitations

There are several potential limitations to this work, which we describe and address more thoroughly in Supplemental Information. Here we provide a brief description of some of the limits of our results:

#### Choice of PSM scoring algorithm

PMD-FDR may have more limited value when used on results from algorithms (e.g. MaxQuant) that explicitly use PMD in their scoring of PSMs. To investigate this question, we applied MaxQuant to the *Pyrococcus* dataset and found that although PMD-FDR has a less pronounced effect in distinguishing high scoring but most likely false PSMs from true hits, we were still able to distinguish these populations based on PMD. Because of the heavy weighting by MaxQuant to results with low PMD, rejecting the false matches came at the cost of also discarding a high proportion of true matches. Supporting Information provides a detailed description of these results.

A related possible limitation from the scoring algorithm is insufficient identification of Good hits, which could lead to over-confidence in results. This algorithm requires a large list of highly confident, correct hits to sufficiently describe the distribution of true-hits. The number of Good PSMs should be in the thousands. This problem can arise either from the choice of PSM scoring algorithm or from an implicit bias in the experiment itself. Similarly, insufficient identification of Bad hits could lead to *under*-confidence in results. If there are a large number of bad hits with correct PMD, the distribution of bad hits will be skewed towards increased FDR scores for items with the correct PMD.

#### Experimental and data acquisition issues

If we have no good data (for example, there is no protein in the sample), the definition of “Good” becomes impossible to define. However, viewing the distribution of PMDs makes this quite apparent – instead of a combination of a flat and peaked distribution, you simply see a flat distribution or a peak that is similar to that of the decoy data.

Other inherent characteristics of detected peptides and MS/MS acquisition may also be limiting. Deamidation can be incorrectly assigned to an isotopic peptide, leading to an approximately 5-ppm discrepancy. These PSMs have the correct sequence but are incorrectly classified as an isotopic peak. PMD-FDR will classify them as probably incorrect if the instrument has better than 5-ppm resolution. Similar problems could arise for other small mass discrepancies. Additionally, chimeric spectra will be problematic for PMD-FDR to the extent that multiple peptides have different precursor masses. Co-eluted peptides with identical chemical compositions (identical numbers of each element) will have the same properties as a singly-eluted peptide with regards to PMD-FDR but that will not be the case for co-eluted peptides with different chemical compositions – the measured precursor mass will be altered by the co-elution, making the PMD an untrustworthy measure for that PSM.

### PMD Algorithm

Correctness, according to the PMD-FDR calculation, is entirely determined by precursor mass. It can suggest exclusion but not inclusion – a high-scoring peptide with a large PMD-FDR should be treated with skepticism, but a peptide with a small PMD-FDR should not be included on the basis of the PMD-FDR score alone. A good example of this latter case is a PSM with small PMD from a MS/MS spectrum for which we have no other good evidence, such as some level of annotated fragmentation peaks consistent with the putative peptide sequence and a reasonably high PSM score assignment from this initial algorithm.

### Applications and Potential extensions

Our PMD framework for identifying high-scoring PSMs that are likely to be false-hits should have immediate application in contemporary shotgun proteomics. One of our motivations in developing this framework was to provide a post-analysis tool for rigorously assessing accuracy of PSMs/peptides of interest to researchers conducting proteogenomic or metaproteomic studies. For example, proteogenomics studies match MS/MS to sequence databases containing both novel variant sequences and reference sequences^4–5^. Researchers rely heavily on scores assigned to single PSMs matching variant sequences to determine whether these are worth further empirical examination. Our PMD-based framework provides an additional assessment of the accuracy of any given PSM, providing researchers an automated means to prioritize those matches to variant sequences most likely to be correct, and therefore of highest priority for further validation (e.g. confirmation with synthetic peptides, development of targeted MS-based methods etc.). Those PSMs flagged as potentially false by PMD-FDR scoring can be dealt with at the discretion of the researcher – potentially rejected or subjected to further scrutiny to validate their veracity. In addition to its immediate value, a number of future avenues can be pursued to extend its functionality and increase its utility, including implementation within existing workflows (e.g. see^5, 20^). We outline several of these in the Supplemental Information.

### Conclusions

Our goal was to create a post-analysis tool that would allow us to automate the identification of high-scoring PSMs which are more likely false-hits, focusing on PSM-centric applications such as proteogenomics and metaproteomics. We selected PMD as our primary input because it was universally available and because, intuitively, true- and false-hit distributions for PMD should be quite distinctive. Along the way we found that all of our assumptions about PMD distributions were incorrect; in particular, these distributions were not fixed within an LC-MS/MS acquisition, let alone across separate LC-MS/MS runs.

By addressing these issues and finding data features that *are* invariant across an experiment, we have created the PMD-FDR measure. By rigorously testing assumptions underlying other PMD-based methods, we have implemented a method to filter PSMs that complements the upstream scoring of PSMs by conventional sequence database searching programs. As such, our PMD-FDR method is agnostic to the sequence database-searching program used and provides a means to assess accuracy of PSMs and the confidence assigned to them by these programs. This methodology should find wide applicability in contemporary shotgun proteomics workflows, especially in applications that depend heavily on PSM accuracy, such as the identification of PTMs, variant sequences in proteogenomics, or species- or isoform-specific peptides detected using metaproteomics.

### Software Availability

The software used to generate the figures in this paper has been released on GitHub: https://github.com/slhubler/PMD-FDR-for-paper

The PMD-FDR algorithm has been implemented as a Galaxy tool, and is available in the Galaxy Tool Shed (https://toolshed.g2.bx.psu.edu/view/galaxyp/pmd_fdr/5cc0c32d05a2) for public use. The link to the Galaxy tool development repository is here: https://github.com/galaxyproteomics/tools-galaxyp/tree/master/tools/pmd_fdr. Documentation on deploying tools from the Tool Shed in any Galaxy instance can be found here: https://galaxyproject.org/admin/tools/add-tool-from-toolshed-tutorial/.

A monolithic version of the software, designed for use as a simple Galaxy-P^5, 20^ module is also on GitHub: https://github.com/slhubler/PMD-FDR-for-Galaxy-P

## Supporting information

Supporting information for main text

## Supporting Information

The following supporting information is available free of charge at ACS website http://pubs.acs.org

Supporting Data S1: A detailed description of results from application of PMD-FDR on representative datasets is provided, along with additional details on limits and conditions for applying the algorithm, including analysis of results from MaxQuant.

## Acknowledgements

We extend our gratitude to Marc Vaudel and Harald Barsnes for discussions on PeptideShaker confidence scoring. We thank Colleen Hayes for assistance with figure design. This work was funded in part by NSF award 1458524 and NIH award U24CA199347 to T.J. Griffin and the Galaxy for proteomics (Galaxy-P) research team.

## Conflict of Interest Disclosure

The authors declare no competing financial interest.

## References

1. Abraham, P. E.; Giannone, R. J.; Xiong, W.; Hettich, R. L., Metaproteomics: extracting and mining proteome information to characterize metabolic activities in microbial communities. Current protocols in bioinformatics 2014, 46, 13.26.1–14.

2. Muth, T.; Renard, B. Y.; Martens, L., Metaproteomic data analysis at a glance: advances in computational microbial community proteomics. Expert review of proteomics 2016, 13 (8), 757–69.

3. Ruggles, K. V.; Krug, K.; Wang, X.; Clauser, K. R.; Wang, J.; Payne, S. H.; Fenyo, D.; Zhang, B.; Mani, D. R., Methods, Tools and Current Perspectives in Proteogenomics. Molecular & cellular proteomics : MCP 2017, 16 (6), 959–981.

4. Nesvizhskii, A. I., Proteogenomics: concepts, applications and computational strategies. Nature methods 2014, 11 (11), 1114–25.

5. Chambers, M. C.; Jagtap, P. D.; Johnson, J. E.; McGowan, T.; Kumar, P.; Onsongo, G.; Guerrero, C. R.; Barsnes, H.; Vaudel, M.; Martens, L.; Gruning, B.; Cooke, I. R.; Heydarian, M.; Reddy, K. L.; Griffin, T. J., An Accessible Proteogenomics Informatics Resource for Cancer Researchers. Cancer Res 2017, 77 (21), e43–e46.

6. Heyer, R.; Schallert, K.; Zoun, R.; Becher, B.; Saake, G.; Benndorf, D., Challenges and perspectives of metaproteomic data analysis. Journal of Biotechnology 2017, 261, 24–36.

7. Shteynberg, D.; Nesvizhskii, A. I.; Moritz, R. L.; Deutsch, E. W., Combining results of multiple search engines in proteomics. Molecular & cellular proteomics : MCP 2013, 12 (9), 2383–93.

8. Burger, T., Gentle Introduction to the Statistical Foundations of False Discovery Rate in Quantitative Proteomics. J Proteome Res 2018, 17 (1), 12–22.

9. Tang, W. H.; Shilov, I. V.; Seymour, S. L., Nonlinear Fitting Method for Determining Local False Discovery Rates from Decoy Database Searches. Journal of Proteome Research 2008, 7 (9), 3661–3667.

10. Eng, J. K.; Searle, B. C.; Clauser, K. R.; Tabb, D. L., A face in the crowd: recognizing peptides through database search. Molecular & cellular proteomics : MCP 2011, 10 (11), R111.009522.

11. Wenger, C. D.; Phanstiel, D. H.; Lee, M. V.; Bailey, D. J.; Coon, J. J., COMPASS: a suite of pre- and post-search proteomics software tools for OMSSA. Proteomics 2011, 11 (6), 1064–1074.

12. Cox, J.; Michalski, A.; Mann, M., Software Lock Mass by Two-Dimensional Minimization of Peptide Mass Errors. Journal of The American Society for Mass Spectrometry 2011, 22 (8), 1373–1380.

13. Petyuk, V. A.; Mayampurath, A. M.; Monroe, M. E.; Polpitiya, A. D.; Purvine, S. O.; Anderson, G. A.; Camp, D. G.; Smith, R. D., DtaRefinery, a Software Tool for Elimination of Systematic Errors from Parent Ion Mass Measurements in Tandem Mass Spectra Data Sets. Molecular & cellular proteomics : MCP 2010, 9 (3), 486–496.

14. Epstein, J. A.; Blank, P. S.; Searle, B. C.; Catlin, A. D.; Cologna, S. M.; Olson, M. T.; Backlund, P. S.; Coorssen, J. R.; Yergey, A. L., ProteinProcessor: A probabilistic analysis using mass accuracy and the MS spectrum. Proteomics 2016, 16 (18), 2480–90.

15. Keller, A.; Nesvizhskii, A. I.; Kolker, E.; Aebersold, R., Empirical Statistical Model To Estimate the Accuracy of Peptide Identifications Made by MS/MS and Database Search. Analytical Chemistry 2002, 74 (20), 5383–5392.

16. Käll, L.; Storey, J. D.; MacCoss, M. J.; Noble, W. S., Posterior Error Probabilities and False Discovery Rates: Two Sides of the Same Coin. Journal of Proteome Research 2008, 7 (1), 40–44.

17. Lippencott, J. Question Assumptions. https://proteomesoftware.zendesk.com/hc/enus/articles/115002213066-Question-Assumptions (accessed April 5, 2019).

18. Searle, B. C., Scaffold: a bioinformatic tool for validating MS/MS-based proteomic studies. Proteomics 2010, 10 (6), 1265–9.

19. Shilov, I. V.; Seymour, S. L.; Patel, A. A.; Loboda, A.; Tang, W. H.; Keating, S. P.; Hunter, C. L.; Nuwaysir, L. M.; Schaeffer, D. A., The Paragon Algorithm, a next generation search engine that uses sequence temperature values and feature probabilities to identify peptides from tandem mass spectra. Molecular & cellular proteomics : MCP 2007, 6 (9), 1638–55.

20. Blank, C.; Easterly, C.; Gruening, B.; Johnson, J.; Kolmeder, C. A.; Kumar, P.; May, D.; Mehta, S.; Mesuere, B.; Brown, Z.; Elias, J. E.; Hervey, W. J.; McGowan, T.; Muth, T.; Nunn, B.; Rudney, J.; Tanca, A.; Griffin, T. J.; Jagtap, P. D., Disseminating Metaproteomic Informatics Capabilities and Knowledge Using the Galaxy-P Framework. Proteomes 2018, 6 (1), 7.

21. Cesnik, A. J.; Shortreed, M. R.; Sheynkman, G. M.; Frey, B. L.; Smith, L. M., Human Proteomic Variation Revealed by Combining RNA-Seq Proteogenomics and Global Post-Translational Modification (G-PTM) Search Strategy. J Proteome Res 2016, 15 (3), 800–8.

22. Heydarian, M.; Luperchio, T. R.; Cutler, J.; Mitchell, C. J.; Kim, M.-S.; Pandey, A.; Sollner-Webb, B.; Reddy, K., Prediction of Gene Activity in Early B Cell Development Based on an Integrative Multi-Omics Analysis. Journal of proteomics & bioinformatics 2014, 7, 1000302.

23. Rudney, J. D.; Jagtap, P. D.; Reilly, C. S.; Chen, R.; Markowski, T. W.; Higgins, L.; Johnson, J. E.; Griffin, T. J., Protein relative abundance patterns associated with sucrose-induced dysbiosis are conserved across taxonomically diverse oral microcosm biofilm models of dental caries. Microbiome 2015, 3, 69–69.

24. Wong, C. C.; Cociorva, D.; Miller, C. A.; Schmidt, A.; Monell, C.; Aebersold, R.; Yates, J. R., 3rd, Proteomics of Pyrococcus furiosus (Pfu): Identification of Extracted Proteins by Three Independent Methods. J Proteome Res 2013, 12 (2), 763–70.

25. Vaudel, M.; Burkhart, J. M.; Breiter, D.; Zahedi, R. P.; Sickmann, A.; Martens, L., A complex standard for protein identification, designed by evolution. J Proteome Res 2012, 11 (10), 5065–71.

26. Jagtap, P.; Goslinga, J.; Kooren, J. A.; McGowan, T.; Wroblewski, M. S.; Seymour, S. L.; Griffin, T. J., A two-step database search method improves sensitivity in peptide sequence matches for metaproteomics and proteogenomics studies. Proteomics 2013, 13 (8), 1352–7.

27. Barsnes, H.; Vaudel, M., SearchGUI: A Highly Adaptable Common Interface for Proteomics Search and de Novo Engines. J Proteome Res 2018, 17 (7), 2552–2555.

28. Vaudel, M.; Burkhart, J. M.; Zahedi, R. P.; Oveland, E.; Berven, F. S.; Sickmann, A.; Martens, L.; Barsnes, H., PeptideShaker enables reanalysis of MS-derived proteomics data sets. Nature Biotechnology 2015, 33, 22.

29. R Core Team R: A language and environment for statistical computing. R Foundation for Statistical Computing. http://www.R-project.org/.

30. Nesvizhskii, A. I., A survey of computational methods and error rate estimation procedures for peptide and protein identification in shotgun proteomics. Journal of proteomics 2010, 73 (11), 2092–2123.

