## Supporting information for main text for "Challenges in Peptide-Spectrum Matching: a Robust and Reproducible Statistical Framework for Removing Low-Accuracy, High-Scoring Hits"

**Table of Contents**

Application to Datasets____S-2

Limitations____S-6

Application to results from an algorithm explicitly using PMD in its scoring: MaxQuant____S-11

Bibliography____S-14

### Application to datasets

In the main text we focused on results for a single dataset, Oral 737 (two-step). However, we intend this tool to be applied to other datasets and project types. We performed similar analyses for three other datasets to show that the general findings (PMD-Shift, Decoy-Mode, and Decoy-Tails, and invariance of the modified PMD) hold for these datasets. In addition, each new analysis offers an additional insight into the dataset or PMD analysis in general.

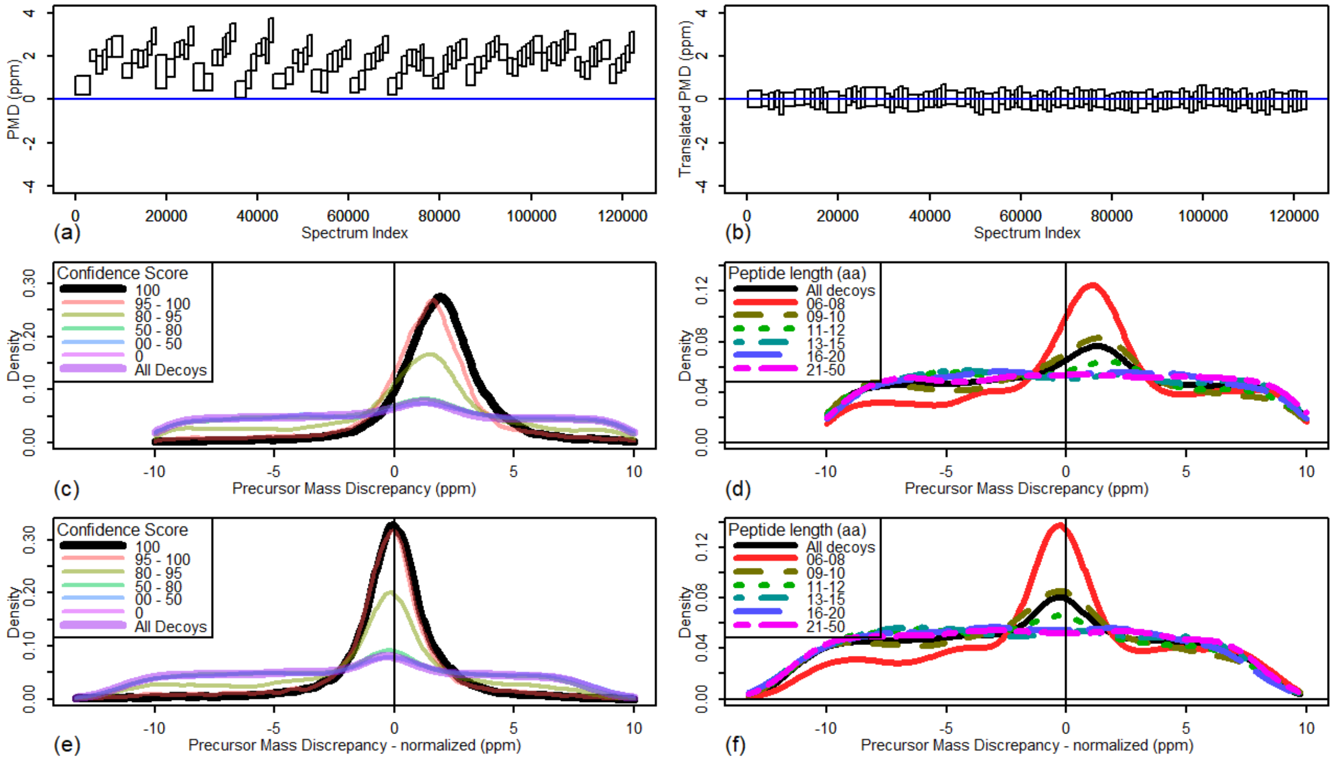


Figure S- 1. Analysis of Oral 737 NS – two-step. Middle 25% Precursor Mass Discrepancy (PMD), good-testing data only, binned into 100 sets by spectrum index, (a) before and (b) after PMD translation. Distribution of PMD, good-testing data only, by score, (c) before and (e) after PMD translation. Distribution of PMD, decoy data only, by peptide length, (d) before and (f) after PMD translation.

For the sake of comparison, we include a single figure (Figure S- 1) representing most of the Oral_737_NS_two_step analysis. Of particular note is that this dataset is a metaproteomics dataset. Of the four datasets analyzed, this one had the most pronounced Decoy-Mode. As shown in the main text, all of the features of interest are present in this figure.

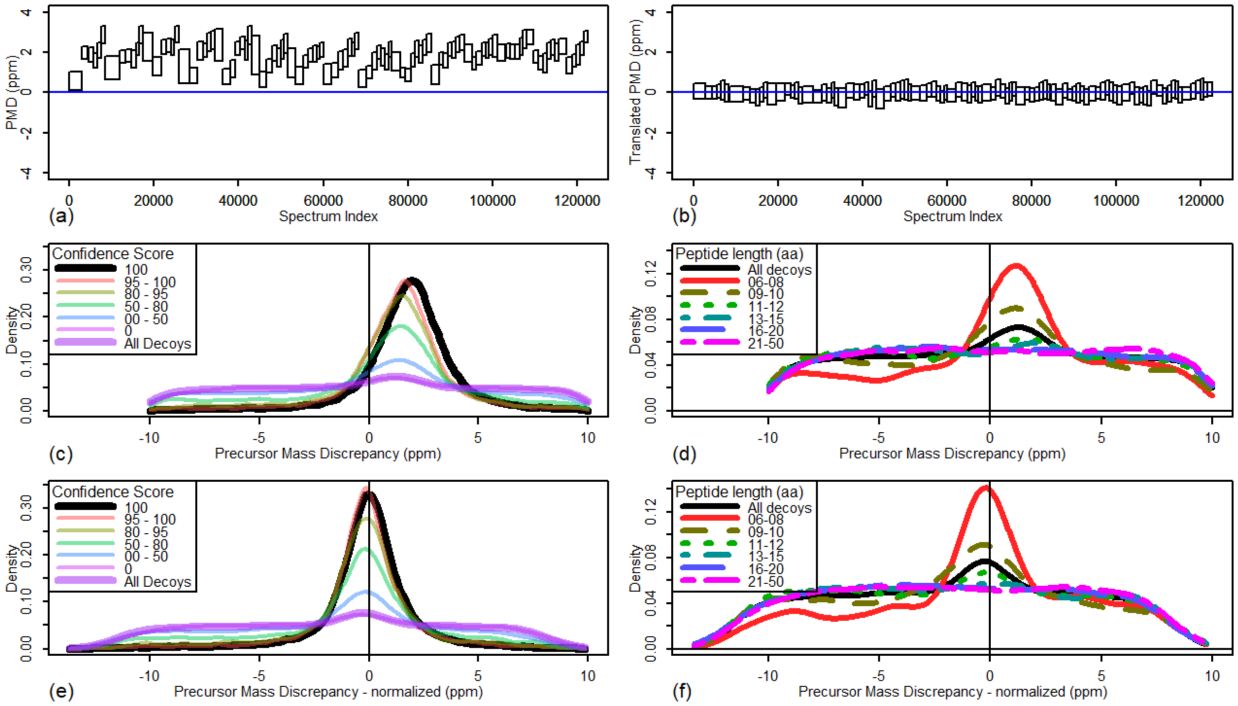


Figure S- 2. Analysis of Oral 737 NS – combined. Middle 25% Precursor Mass Discrepancy (PMD), good-testing data only, binned into 100 sets by spectrum index, (a) before and (b) after PMD translation. Distribution of PMD, good-testing data only, by score, (c) before and (e) after PMD translation. Distribution of PMD, decoy data only, by peptide length, (d) before and (f) after PMD translation.

We included (Figure S- 2) to show similarities of PMD-FDR analyses even when the underlying FDR methodologies were different – the two datasets used different metaproteomics workflows, applied to the same spectra. Note that, while there are some differences (in particular, there were more high-scoring PSMs in the second workflow (combined) than in the first), they are qualitatively the same. In other words, the estimation of the True-Hit and False-Hit distributions are nearly identical and, therefore, so would the local FDR values be nearly identical for a given PSM.

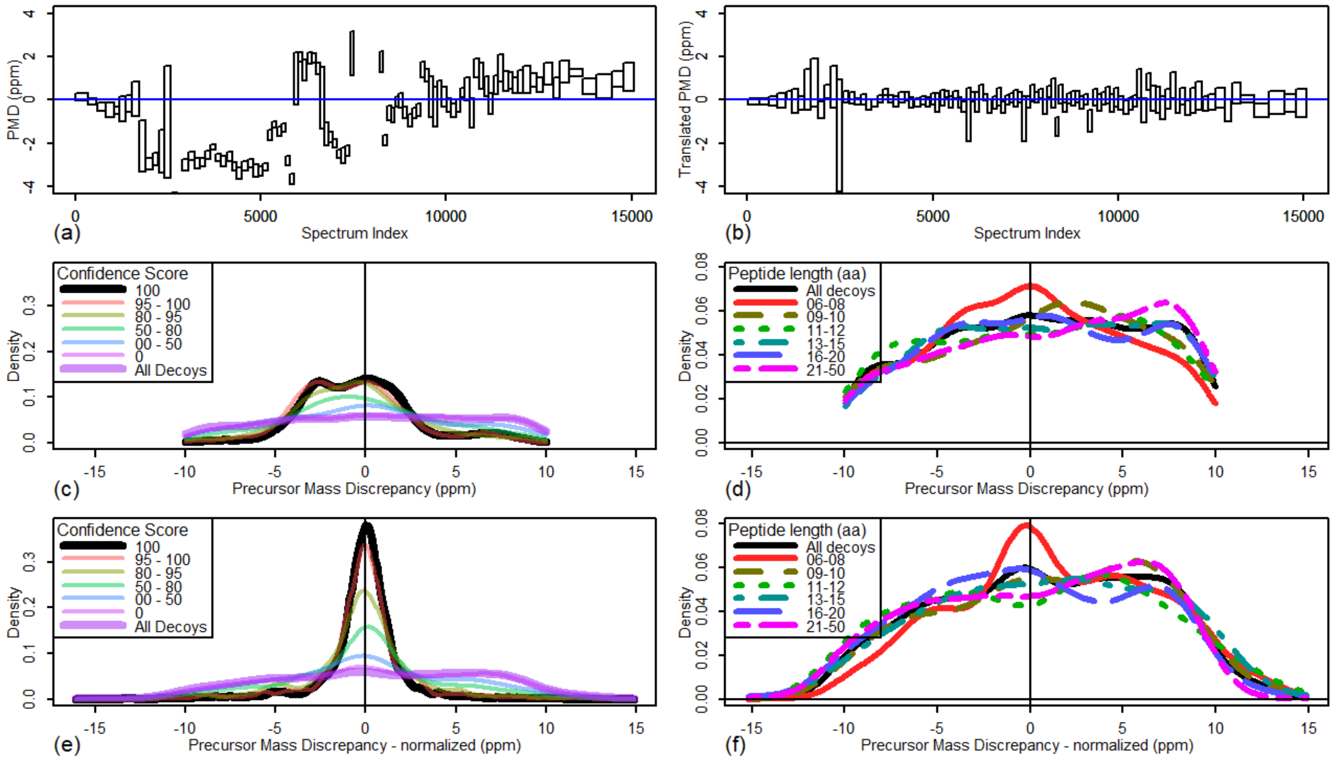


Figure S- 3. Analysis of Pyrococcus dataset. Middle 25% Precursor Mass Discrepancy (PMD), good-testing data only, binned into 100 sets by spectrum index, (a) before and (b) after PMD translation. Distribution of PMD, good-testing data only, by score, (c) before and (e) after PMD translation. Distribution of PMD, decoy data only, by peptide length, (d) before and (f) after PMD translation.

We included the Pyrococcus_tr data (Figure S- 3) as an example of a small, traditional experiment – it included only one file's worth of spectra and represented a relatively simple workflow, although because the human genome was included in the analysis of a single, small proteome, most of the reference database was not relevant. Nonetheless, we were able to identify several useful features from this dataset. In particular, this dataset had large, sudden PMD shifts (as much as 11 ppm). This created a multi-modal PMD distribution. Ironically, this dataset may have come from the most precise instrument (Figure S- 3e), although it was certainly the least accurate (Figure S- 3c), as observed by the width of the distribution represented by the black curves.

Interestingly, the Decoy-Mode was only noticeable *after* translation and only for the smallest peptides (less than 9 amino acids). This is probably due, though, to the total number of spectra being one-twentieth as large as that in other experiments.

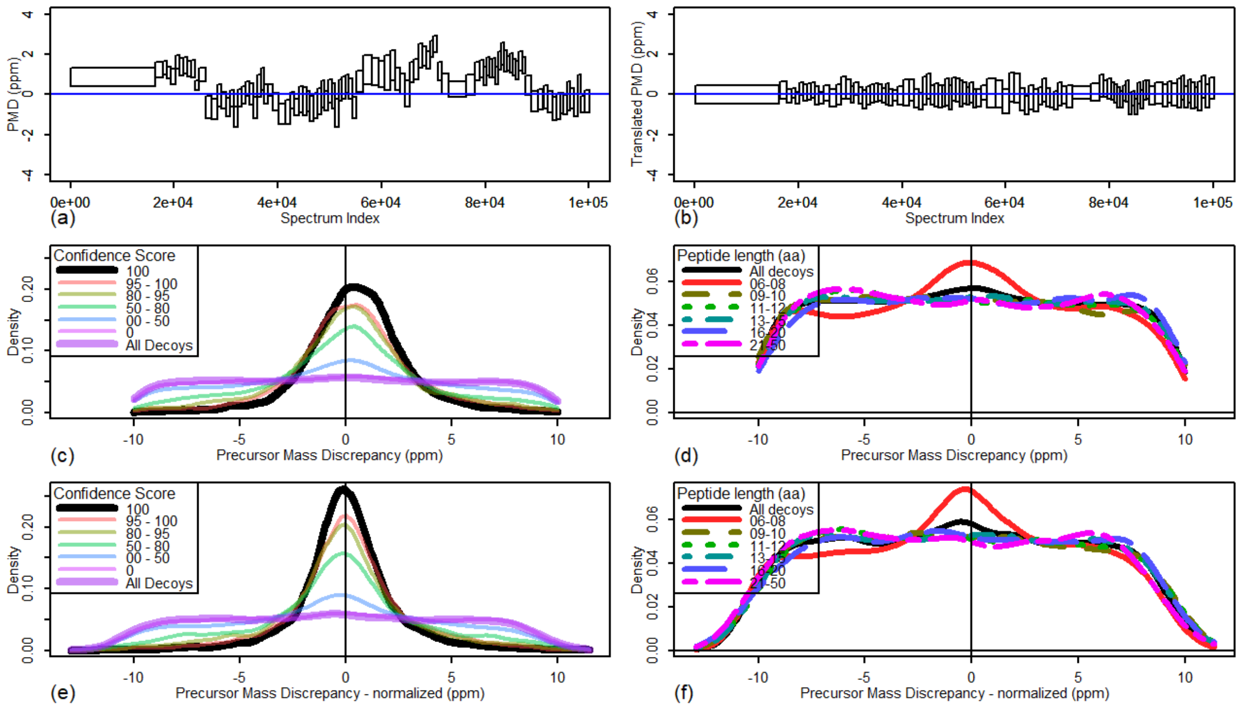


Figure S- 4. Analysis of Mouse proteogenomic dataset. Middle 25% Precursor Mass Discrepancy (PMD), good-testing data only, binned into 100 sets by spectrum index, (a) before and (b) after

PMD translation. Distribution of PMD, good-testing data only, by score, (c) before and (e) after PMD translation. Distribution of PMD, decoy data only, by peptide length, (d) before and (f) after PMD translation.

Finally, we included a dataset produced from a proteogenomics workflow (Figure S- 4). Other than the fact that this dataset appears to have the lowest precision, it is very similar to the other analyses. Note, in particular that all four datasets have the features described earlier.

### Limitations

Within the main text we have summarized some of the limitations of the approach. Here we expand on these in more detail.

Calculating PMD-FDR described in this work requires several features working correctly. Below we describe what can go wrong and should be considered when applying this method.

*Explicit use of PMD in scoring*

In theory, we assume that the scoring mechanism does not explicitly depend on PMD; if it does then the score will be a function of PMD, which could potentially bias PMD-FDR results. As mentioned in the introduction, remarkably few algorithms explicitly depend on PMD, though those few that do are widely used (such as Percolator[1]). We have not yet investigated what can go wrong if one uses such a scoring mechanism. However, because we use empirical distributions to define the good- and bad-hit distributions, the algorithms should still work; we suspect that the algorithm will report a bias for good- and bad- hits towards zero PMD. Since that bias will exist for both groups, we should still be able to calculate iFDR and gFDR; the catch will be that these distributions should also be constant throughout the experiment, independent of other variables, a condition that may not exist in this case. Note that this will probably report a higher FDR than the original algorithm since the bias towards a zero PMD will exist for all spectra, including decoys.

It is important to note that this algorithm will not be sensitive to the nearly universal practice of restricting peptide candidates based on their difference from the precursor mass (i.e. the PMD); this is an *implicit* use of PMD that does not directly affect the score of a PSM, merely which scores we test. Neither will our algorithm be harmed by a PMD renormalization, such as used by Perseus[2], although this normalization may make the PMD-translation part of our algorithm redundant. The critical information that the algorithm needs is that there are enough examples of "good" and "bad" hits to provide good estimates of those distributions. Thus, it is important to include most or all PMDs, not just those with high confidence. Including only the 1% TD-FDR hits, for example, will cause difficulties since there will not be enough decoys to create a good distribution of the bad hits, especially those with a poor PMD.

*When "good" is not good enough*

The technique of using PMD to create a PMD-FDR requires an estimate of the true-hits generated from the distribution of "good" hits (or, perhaps, "good enough" hits). However, if there are a significant number of false hits in the distribution, the distribution will spread, artificially inflating our confidence in PSMs.

Note that this can be detected qualitatively by looking at how the distribution tails off – if the tails of the estimated true hits are not close to zero at the edges of the region of consideration, then either the PMD restriction region is too small or there are too many false hits in the good list.

*When "bad" is not bad enough*

The mirror image of the "not good enough" issue is the "not bad enough" issue. If true hits exist in the "bad" group then we are not correctly estimating the false hit distribution, leading to a deflation of our confidence in PSMs.

Note that we have already dealt with this issue in a different guise: the Decoy-Mode exists because the smaller random peptides are more likely to have the correct mass (though, presumably, not the correct sequence). This is an important point – PMD-FDR refers to the probability that the PSMs peptide has an incorrect mass; it says nothing about being the incorrect sequence if the mass is accurate. It is true that a correct identification must have a correct mass but the converse is not true except in cases where only one option is possible.

We can detect this by noting a Decoy-Mode; if we find two bad populations with slightly different density functions then one probably contains PSMs with the correct precursor mass, confounding our results. In this sense large-decoy peptides are "worse" than small-decoy peptides.

Note that the Decoy-Mode reported in this work probably arises from short (decoy) peptides having exactly the same precursor mass as the correct peptide (i.e., the random peptide has exactly the same number of N, O, H, C, and S atoms as the true peptide). This is much more likely for small peptides than for large ones: the number of distinct peptigenic chemical masses made of five elements (N, O, H, C, S) within a given range (say, within 1 Da) of a given mass grows as a 5$^{th}$ power of mass[3]. Thus, the probability that a random peptide within a fixed size window of the measured mass has exactly the same mass as the true peptide is roughly proportional to the inverse of the 5$^{th}$ power of mass/number of amino acids. Since we are working with a window size that increases linearly with the size of the peptide (the units of accuracy in our cases is ppm instead of Da), this random peptide is proportional to the inverse of the 4$^{th}$ power of mass / number of amino acids. In particular, if we stretch this approximation, possibly to its breaking point, we find that going from a length of 6 amino acids to 12 amino acids (a doubling) will change the probability of an exact mass match by a random peptide by a factor of $2^{-4} = 1/16$ . So, for example, if 64% of random peptides of length 6 have the correct precursor mass (which they appear to have in the first dataset), we would expect 4% of random peptides of length 12 to have the correct precursor mass.

*When scores go wrong*

For this project, we used PeptideShaker's Confidence score as the initial score of a PSM, allowing us to select good spectra, those that were very likely to be true hits. In fact, any PSM scoring algorithm should work, as long as we can identify such a group of likely true hits. Ideally, for example, it is a group that naturally excludes decoys, as Confidence=100 does. However, if the score is not able to identify a select group of PSMs that are nearly all true hits then this algorithm, as it stands, cannot function properly. That is the price we pay for using good hits to estimate true hits. It may be possible to derive the PMD distribution of true- and false-hits by using expectation-maximization or similar techniques but our initial forays into this line of reasoning led to poor performance, both in quality and in resource use.

*Chimeric spectra*

Chimeric spectra provide a particular challenge for this algorithm – a correctly identified peptide in a chimeric spectrum may or may not have an accurate precursor mass, possibly being the combination of two or more distinct masses. If one uses an algorithm to deconvolute multiple peptides, one would also need to deconvolute the precursor masses in order to correctly assess PMD. If that is possible then PMD-FDR should work; otherwise every chimeric spectrum is effectively a noise spectrum with regards to this algorithm, a type of noise that has not been modeled here. In other words, further investigation would be required to determine if this algorithm would work with chimeric spectra and associated deconvolution algorithms.

*Deamidation*

One interesting side note is that, because we are using empirical distributions, we found that deamidated peptides could be discovered without search by looking for peptides that were misclassified as being isotopes and had a corrected PMD of approximately -5 ppm (depending on the actual peptide size). The actual difference between a deamidated peptide and its isotopic cousin (assuming $C^{13}$ isotope) is 0.00447 Da; thus, a -5 ppm difference corresponds to peptides of approximately 1000 Da or 10 amino acids. We could easily identify the deamidated peptides by plotting two datasets against each other, one with deamidation (and no isotopes) and the other without deamidation (and with isotopes).

*Correct peptide or correct mass?*

TD-FDR estimates the probability that the precursor mass of the peptide reported for a spectrum *does not match* the precursor mass of the actual peptide in the sample. In other words, PMD-FDR represents the likelihood that the reported peptide has the wrong mass and, therefore, is the wrong peptide. It is important to note that this measure does not have a contrapositive: an accurate mass does not improve the probability that we have identified the correct peptide, at least without knowing what other peptides could have the same mass. In other words, it is an *optimistic* measure of FDR, giving us more confidence in a PSM than may be entirely warranted. However, since it is empirically less optimistic than other, similar, measures, we do not consider this to be a problem at present; it is being used to report on cases where the precursor mass does not appear to be correct.

*Calculating credible intervals*

While our primary analysis did not use credible intervals, we did use it to create Figure 7 of the main text, where we calculated the 95% credible interval of a proportion. A *credible* interval differs from a confidence interval, approaching a question from a Bayesian perspective instead of a frequentist. It is useful for identifying the most likely range of values for a parameter (proportion, in our case), rather than the range of values that are likely to be produced given that you have an estimate of the mean (the standard confidence interval). For our calculation we chose to assume a uniform prior and approximate the highest posterior density interval. This choice has three useful properties: 1) it is guaranteed to include the observed proportion; 2) it is the smallest possible credible interval of a given confidence; and 3) it is the decision theoretic optimal interval[16]. We estimated the interval by first assuming a uniform prior, dividing the interval [0,1] into 1001 values and calculating the probability, for each, that it would produce the observed data (the conditional probability that the value was produced given that the individual probability had the value in question). Next, we divided each by the sum of the conditional probabilities; we now have an estimate of the probability that the value produced the data (given a uniform prior). Finally, we sort the values (largest to smallest), adding them together until the sum added to 95% or more; this is an estimate of the 95% credible interval. By using 1001 tests on the interval [0,1], our estimate should be precise to approximately 0.001.

### Application to results from an algorithm explicitly using PMD in its scoring: MaxQuant

We applied PMD-FDR to the MaxQuant analysis of the Pyrococcus data. MaxQuant explicitly utilizes PMD as a criteria for scoring and qualifying identified peptides as being correct. We used evidence.txt as the input file, using the field labeled “Mass error [ppm]” as the PMD. The input score was computed from the following formula:

$$score = 100 \times (1 - PEP)$$

Also, we identified all peptides with a score greater than 99.9999 as "good" peptides identified by MaxQuant; i.e. we changed all of these to 100 so that the algorithm would identify them as good.

It is important to note that evidence.txt is a peptide-centric file and that all values in an outputted "record" are aggregated across several spectra; strictly speaking, this is not consistent with the expectations of PMD-FDR, which was developed for PSM-centric results such as those from commonly used algorithms encapsulated in the SearhGUI/PeptideShaker platform. However, we were able to find analogs in the MaxQuant results compatible with PMD-FDR, with some success.

As expected of an algorithm that renormalizes the PMD, the PMD was centered around zero ppm (Figure S-5). We also see a shift of the "Bad" records towards the center – this was also expected because the algorithm explicitly uses PMD as part of its scoring mechanism. The net result, from the perspective of PMD-FDR is to bias both true and false records toward 0. In reality this is simply a matter of giving more weight to PMD accuracy than is the case in most scoring algorithms.

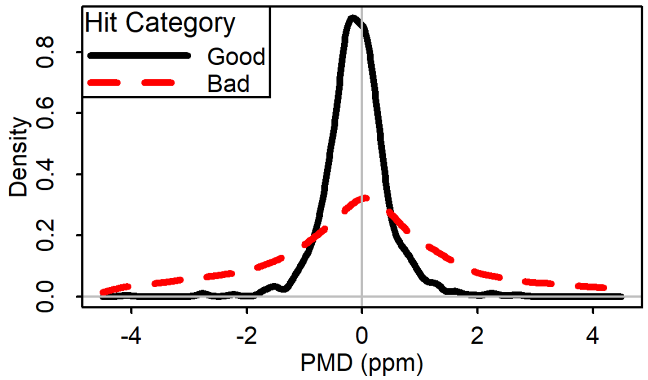


Figure S-5. PMD distribution of peptide identifications from MaxQuant.

Since we applied this analysis to the Pyrococcus data, we were able to see what happens to false records (Figure S-6), indicated both by identifications to human peptide sequences as well as decoys. Interestingly, human-decoy and reverse-decoy had qualitatively identical behaviors –

asymptotically approaching 100% disqualification as we increase the score (i.e. decrease PEP, aka peptide error probability). This disqualification rate on the decoy data is very reassuring – PMD-FDR is doing its job in rejecting high scoring but most likely false results – but it comes at a price: a much higher rejection rate on the peptides that are likely to be correct; for higher scores this rejection rate approaches 50%, as opposed to 10% with PeptideShaker.

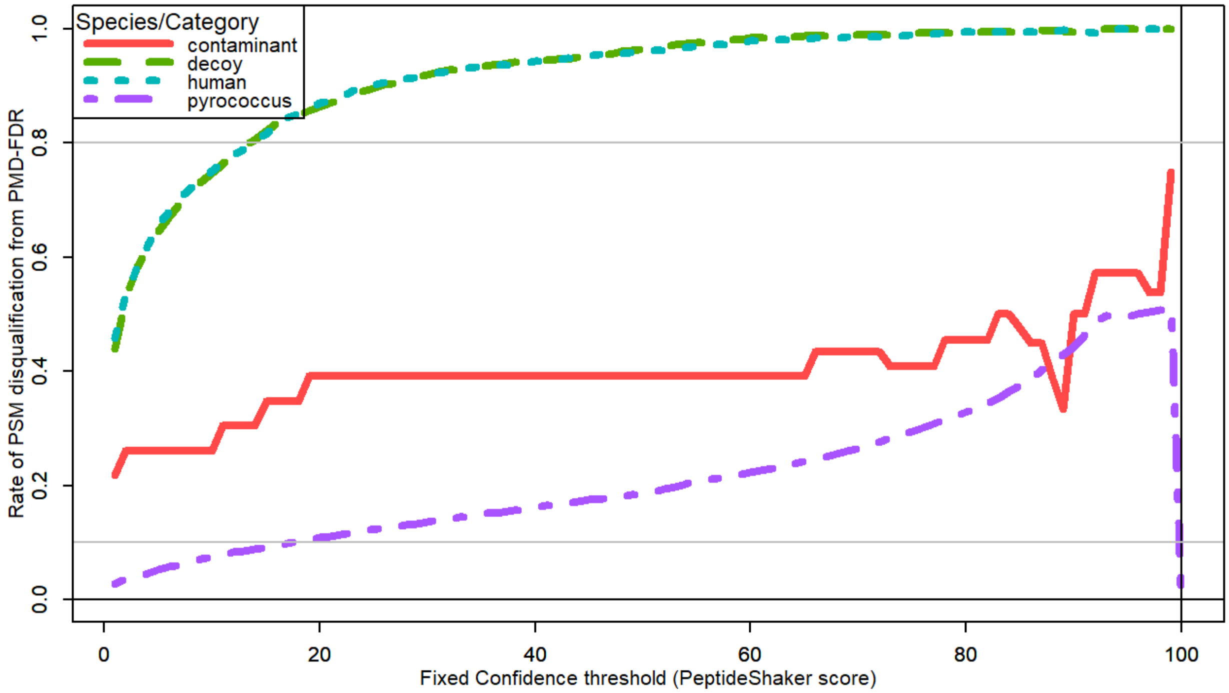


Figure S-6. Rejection rate of all peptides identified by MaxQuant when using PDM-FDR

Finally, we see that the fine structure of the data is unusual when analyzed by PMD-FDR (Figure S-7). In particular, it appears that there is a category of low-scoring peptides (scores of 50-80) centered on +0.5 ppm instead of on zero. This would require investigation but it may be related to our previously discovered misclassification of deamidations as a +1 isotope; because MaxQuant uses PMD as part of its score, such misclassifications would carry a severe score penalty, thus moving them into a lower category. Decoy-mode is also very pronounced (panels d and f in Figure S-7), which is not surprising given that all decoys are selected for low PMD as are matches to forward peptides.  Again, shorter peptide lengths predominate these decoy peptides when analyzed by PMD-FDR.  The longest peptide, however, are not as flatly distributed.

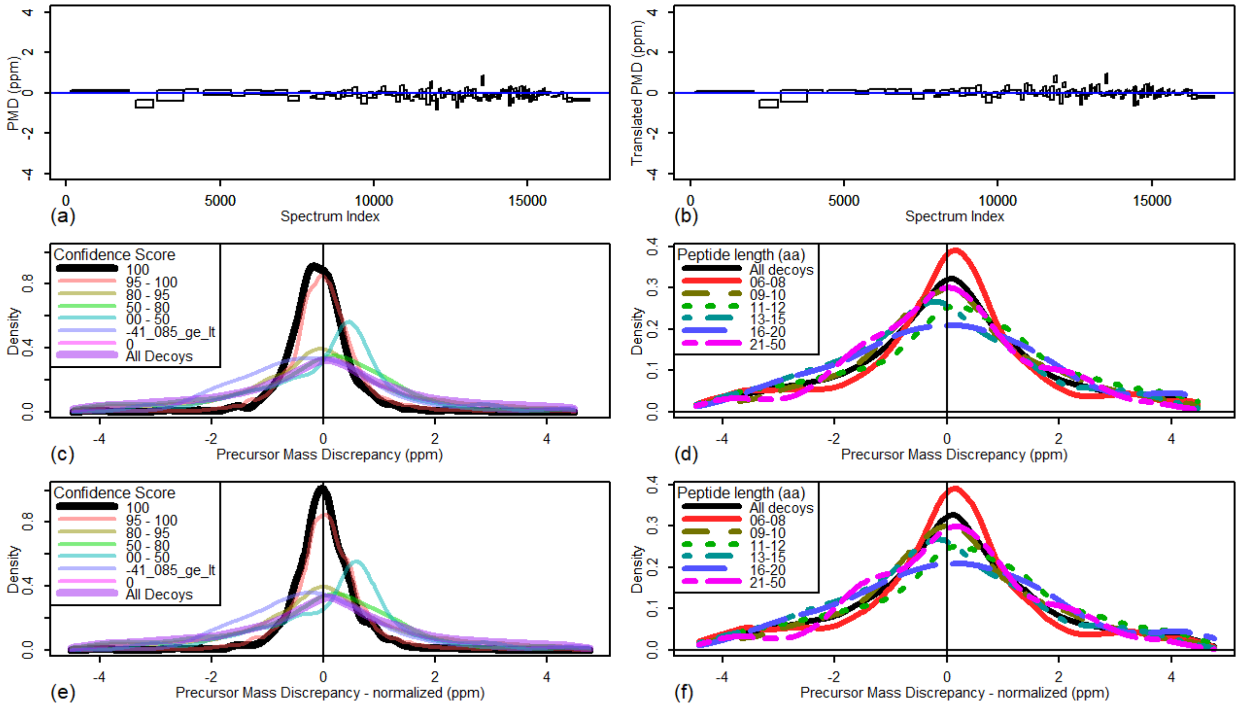


Figure S-7. Analysis of MaxQuant results. Middle 25% Precursor Mass Discrepancy (PMD), good-testing records only, binned into 100 sets by spectrum index, (a) before and (b) after PMD translation. Distribution of PMD, good-testing records only, by score, (c) before and (e) after PMD translation.  Distribution of PMD, decoy data only, by peptide length, (d) before and (f) after PMD translation.

### Bibliography


1. Käll, L.; Storey, J. D.; Noble, W. S., Non-parametric estimation of posterior error probabilities associated with peptides identified by tandem mass spectrometry. *Bioinformatics* **2008,** *24* (16), i42-i48.

2. Cox, J.; Michalski, A.; Mann, M., Software Lock Mass by Two-Dimensional Minimization of Peptide Mass Errors. *Journal of The American Society for Mass Spectrometry* **2011,** *22* (8), 1373-1380.

3. Hubler, S. L.; Craciun, G., Counting chemical compositions using Ehrhart quasi-polynomials. *Journal of Mathematical Chemistry* **2012,** *50* (9), 2446-2470.
